# Interferon-stimulated and metallothionein-expressing macrophages are associated with acute and chronic allograft dysfunction after lung transplantation

**DOI:** 10.1101/2021.06.14.447967

**Authors:** Sajad Moshkelgosha, Allen Duong, Gavin Wilson, Tallulah Andrews, Gregory Berra, Benjamin Renaud-Picard, Mingyao Liu, Shaf Keshavjee, Sonya MacParland, Jonathan Yeung, Tereza Martinu, Stephen Juvet

## Abstract

Lung transplant (LT) recipients experience episodes of immune-mediated acute lung allograft dysfunction (ALAD). ALAD episodes are a risk factor for chronic lung allograft dysfunction (CLAD), the major cause of death after LT. We have applied single-cell RNA sequencing (scRNAseq) to bronchoalveolar lavage (BAL) cells from stable and ALAD patients and to cells from explanted CLAD lung tissue to determine key cellular elements in dysfunctional lung allografts, with a focus on macrophages. We identified two alveolar macrophage (AM) subsets uniquely represented in ALAD. Using pathway analysis and differentially expressed genes, we annotated these as pro-inflammatory interferon-stimulated gene (ISG) and metallothionein-mediated inflammatory (MT) AMs. Functional analysis of an independent set of AMs *in vitro* revealed that ALAD AMs exhibited a higher expression of CXCL10, a marker of ISG AMs, and increased secretion of pro-inflammatory cytokines compared to AMs from stable patients. Using publicly available BAL scRNAseq datasets, we found that ISG and MT AMs are associated with more severe inflammation in COVID-19 patients. Analysis of cells from four explanted CLAD lungs revealed similar macrophage populations. Using a single nucleotide variation calling algorithm, we also demonstrated contributions of donor and recipient cells to all AM subsets early post-transplant, with loss of donor-derived cells over time. Our data reveals extensive heterogeneity among lung macrophages after LT and indicates that specific sub-populations may be associated with allograft dysfunction, raising the possibility that these cells may represent important therapeutic targets.

## Introduction

Lung transplantation (LT) is a life-saving treatment option for advanced lung disease (1), but long-term survival after LT remains relatively poor, despite recent improvements (2). The main factor limiting long-term survival is chronic lung allograft dysfunction (CLAD), a progressive fibrotic process that destroys the lung allograft and is driven by inflammation and alloimmunity. Primary graft dysfunction, acute cellular rejection, antibody-mediated rejection and infection are among the main risk factors for CLAD, but how these insults drive CLAD development remains unknown, because the cellular mechanisms that link these risk factors to organ fibrosis remain incompletely understood.

Bronchoalveolar lavage (BAL) allows sampling of immune cells in the distal pulmonary compartment. Previous BAL cellular phenotyping studies have focused on associations between clinically defined entities (e.g. CLAD) and a limited number of predetermined cell subsets in the BAL (reviewed in (3, 4)), including neutrophils (5) and lymphocytes (6). However, classifying BAL cells in this way underestimates their heterogeneity and overlooks contributions of cell populations not expressing the markers employed. Further, flow cytometric analysis of alveolar macrophages (AMs) – the most abundant cell population in the BAL – is significantly hindered by autofluorescence.

AMs play a critical role in maintaining pulmonary homeostasis through interactions with the alveolar epithelium, and during acute inflammation by orchestrating pro-inflammatory and profibrotic responses through phagocytosis and secretion of inflammatory cytokines and reparative molecules (7-10). The ontogeny of AMs differs significantly between conditions of homeostasis and inflammation. Murine studies suggest that most tissue resident AMs arise during embryogenesis and self-maintain with a minimum contribution from peripheral monocytes (11-13). In contrast, a single-cell RNA sequencing (scRNAseq) study of BAL cells from sex-mismatched LT recipients showed that donor AMs in the human lung are replaced by recipient monocyte-derived macrophages (14). The use of scRNAseq to study AMs has the potential to deepen our knowledge of this cell population by allowing us to uncover important heterogeneity within it.

In recent years, scRNAseq has been applied to the study of lung cells including AM, both in humans and in animal models (14-24). Two recent studies of BAL cells in patients with COVID-19 infection have focused on tracking viral RNA in infected cells (25) and studying immune responses to the disease (26). Both reports revealed substantial differences between AMs from patients with mild disease compared to those with illness. BAL cells have also been studied using scRNAseq in transplant recipients (14) and healthy controls (15). However, to our knowledge, no study has specifically examined the association between disease severity and AM transcriptional states at single cell resolution.

Here, we applied scRNAseq to BAL cells from LT recipients with stable lung function or acute lung allograft dysfunction (ALAD) and to cells from CLAD lung tissue to test the hypothesis that pathogenetically important AM transcriptional states might emerge during lung allograft dysfunction. We observed 14 distinct gene expression programs with evidence for functional specialization of AMs in the BAL; further, two distinct inflammatory macrophage populations seen only in ALAD BAL samples and CLAD lung tissue mirrored similar cellular states in a public dataset of BAL samples obtained from patients with severe COVID-19.

## Materials and Methods

### Ethics statement

The study was approved by University Health Network Research Ethics Board (15-9531). All participants provided written informed consent for sample collection and analyses.

### Study design and patient selection

Six (one female and five male) LT recipients were selected, three with acute lung allograft dysfunction (ALAD, defined as a decrease in the forced expiratory volume in one second [FEV_1_] by 10% or more from the maximum of the two preceding FEV_1_ measurements) and three with stable lung function. Patients with suspected infection (focal opacities on chest imaging and/or mucopurulent secretions at bronchoscopy) were excluded. Demographic characteristics of the study participants are provided in Table 1.

**Table 1.** Clinical characteristics of stable and ALAD patients.

|  | scRNAseq |  | Flow cytometry |  |
| --- | --- | --- | --- | --- |
|  | Stable (n=3) | ALAD (n=3) | Stable (n=4) | ALAD (n=3) |
| Recipient age at LTx (mean) | 63.7±4.7 | 53.7±22.5 | 68.4±3.9 | 71.2±7.5 |
| Male n (%) | 3 (100%) | 2 (66.6%) | 3 (75%) | 2 (66.6%) |
| D/R sex mismatch | 1 (33.3%) | 2 (66.6%) | 2 (50%) | 0 |
| Native lung disease |  |  |  |  |
| Pulmonary Fibrosis/ Interstitial lung dis. | 3 | 0 | 4 | 2 |
| COPD/Emphysma | 0 | 2 | 0 | 0 |
| Cystic Fibrosis | 0 | 1 | 0 | 0 |
| Hypersensitivity Pneumonitis | 0 | 0 | 0 | 1 |
| CMV D/R |  |  |  |  |
| D+/R- | 1 | 1 | 0 | 1 |
| D+/R+, D-/R+ | 2 | 1 | 3 | 2 |
| D-/R- | 0 | 1 | 1 | 0 |
| Previously treated acute cellular rejection | 1 (33.3%) | 0 | 2 (50%) | 1 (33.3%) |
| Respiratory symptoms at time of bronchoscopy | 0 | 1 (33.3%) | 0 | 0 |
| FEV1 at time of bronchoscopy, median (IQR) | 1.9 (1.9, 2.4) | 2.1 (1.7, 2.99) | 2.52 (2.1, 2.7) | 2.2 (1.8, 2.8) |
| Baseline FEV1, median (IQR) | 2.1 (2.09, 2.38) | 2.62 (1.95, 4.31) | 2.66 (2.2, 2.9) | 2.53 (2.1, 3.1) |
| Time from LTx to bronchoscopy (days) | 545.7±181.3 | 821.7±775.9 | 202±151.5 | 211.3±178 |
| Acute rejection grade |  |  |  |  |
| A0 | 3 | 2 | 3 | 2 |
| AX | 0 | 1 | 1 | 1 |
| B-grade |  |  |  |  |
| B0 | 1 | 0 | 1 | 1 |
| BX | 2 | 3 | 3 | 2 |

### BAL sample collection

Patients underwent bronchoscopy for surveillance or for diagnosis of ALAD. BAL was obtained according to a standard protocol in accordance with international guidelines (2). Fresh BAL samples were transferred to the research laboratory on ice within 10 minutes of acquisition.

### scRNAseq sequencing and data analysis

Cells from stable (n=3; 3044 ± 1519 cells) and ALAD (n=3; 2593 ± 904 cells) BAL underwent barcoding and library construction using 10X Genomics 3’ expression V2 chemistry. Library constructs underwent scRNAseq. Data were analysed using packages in R including Seurat for QC, clustering workflow, and sample integration, SingleR for cell annotation, and ClusterProfiler for pathway analysis.

## Results

### Functional diversity of AMs from patients with stable lung function

BAL samples were collected at varying post-LT times from patients who were either stable (n=3) or experiencing ALAD (n=3) (Fig 1A and Table 1). Single cell transcriptomes from each sample were analyzed separately, following standard quality control protocols (Fig S1). We applied a reference-based annotation algorithm (ClusterR) and manual analysis of the top differentially expressed genes to identify cell populations (Fig S2A-C), demonstrating that the samples contained epithelial, endothelial and immune cells; most were AM. Cell-cell interaction analysis using CellChat (27) suggested that interaction amongst AMs are stronger and more complicated than between AMs and other cells (Fig S2D, E). Given the predominance of AM over other cell populations in the BAL samples, we next examined these cells in greater depth.

**Fig 1.**
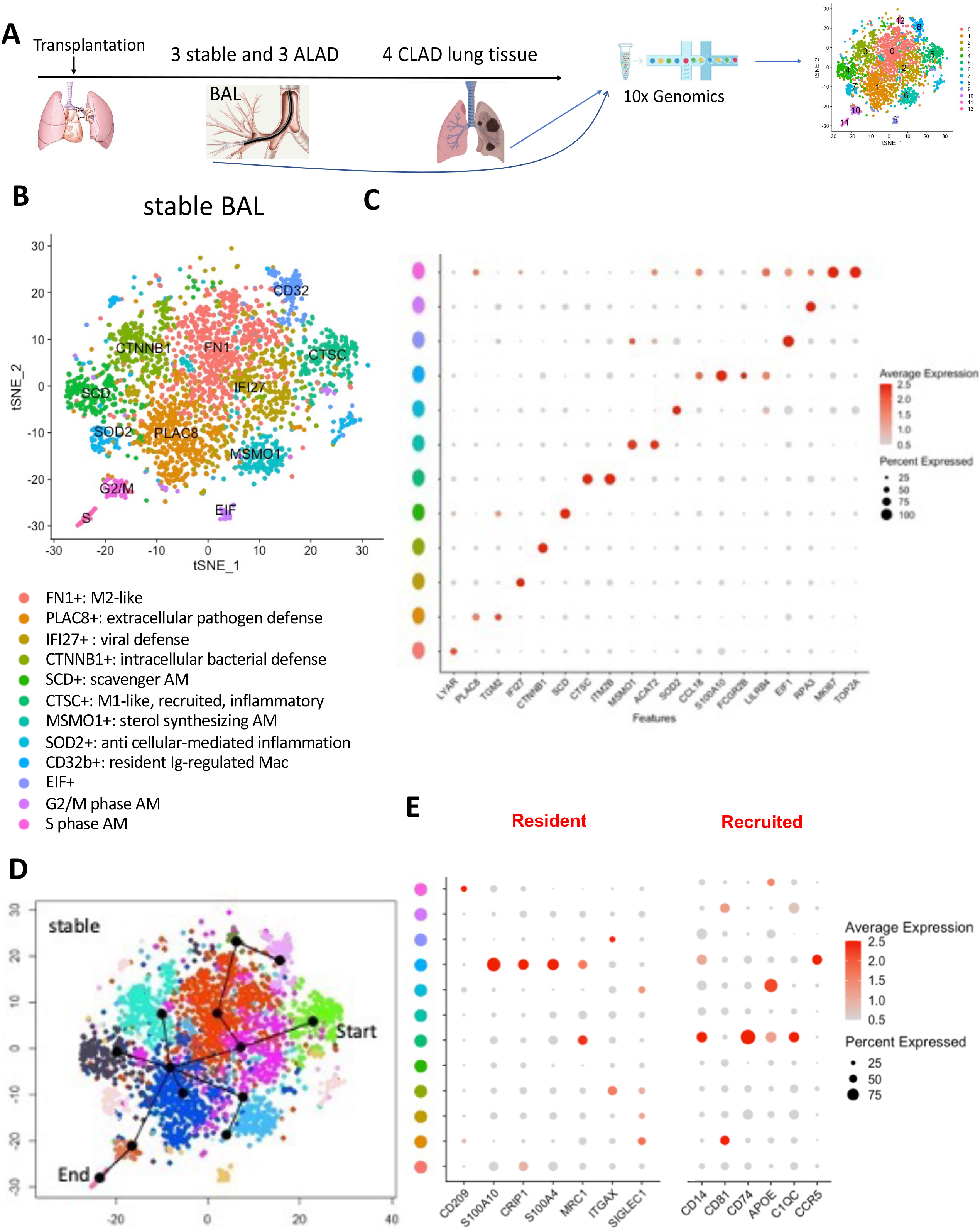
The cellular heterogeneity of AMs from stable LT recipients is revealed by single cell RNA sequencing. **(A)** Study design. Three stable and three ALAD patients with no evidence of pulmonary infection underwent bronchoscopy at varying times post-transplant. CLAD lung tissue (n=4) was obtained at re-transplantation. Cells were subjected to scRNAseq on the 10x genomics platform. **(B)** tSNE plot showing 12 distinct AM populations in BAL from stable patients. Functional annotation of AMs was performed based on top differentially expressed genes and pathway analysis. **(C)** Expression of the top differentially expressed genes in each stable AM cluster (x-axis) according to cluster (y-axis). Circle colour reflects average expression within the cluster, while the size of each circle reflects the percentage of cells within the cluster expressing the indicated gene. **(D)** Pseudotime trajectory analysis was performed using Slingshot. Recently differentiated CD32b^+^ are at the start of pseudotime while cycling AMs are at the end of the trajectory. **(E)** Expression of canonical genes (x-axes) associated with resident (left panel) and recruited (right panel) AMs.

The total number of AMs in the 3 stable BAL samples ranged from 1769 to 4681. We randomly selected a 1000-cell subset (downsample) of AMs from each stable sample to optimize integration. Next, we used SCTransform (28) to integrate scRNAseq data from all three stable samples (Fig S3A) and data from all three ALAD samples (Fig S3B) to minimize batch effects arising from different sequencing runs. Expression of known macrophage transcripts confirmed that the cells were AMs (Fig S4). Clustering revealed 12 subsets (Fig 1B). We then integrated differentially expressed genes and gene set enrichment analysis (GSEA) (Fig S5A) with data reported in the literature to annotate each cluster based on either putative function or cycling status (Fig S5B). We identified three clusters with specialized anti-microbial functions (PLAC8^+^: extracellular pathogen defense; IFI27^+^: viral defense; CTNNB1^+^: intracellular bacterial defense) (29-31), two with specialized immune regulatory functions (SOD2^+^: anti-inflammatory, CD32b^+^: resident Ig-regulated), one with sterol-synthesizing function (MSMO1^+^)(32), one with potential scavenger function (SCD^+^) (33), one with potential healing and/or profibrotic genes (FN1^+^)(34), one with a higher expression of inflammatory markers (CTSC^+^: M1-like, recruited, inflammatory), one with several eukaryotic initiation factors (EIF) (35) and two AM populations in the cell cycle (18, 36).

Of the latter, one had G2/M phase transcripts while the other had S phase transcripts. These data suggest that each AM cluster may have distinct functional properties, some of which have been ascribed to AMs previously (Table 2). Fig 1C shows the most differentially expressed genes that were also expressed in the majority of the cells in each cluster. We also analyzed the incoming and outgoing ligand/receptor signaling network for each cluster (Fig S6) to gain insight into how the clusters may interact with each other.

**Table 2.** Functional annotation of AMs in stable and ALAD BAL samples

| Cluster Id | Cluster specificity/function | Previous similar observations | Reference |
| --- | --- | --- | --- |
| FN1 | M2-like AM | Several top genes (inc. FN1) is found in M2-like cells | Jablonski et al., 2015 (47) |
| PLAC8 | Extracellular pathogen defense AM | Functional importance shown in mouse models | Ledford et al., 2007 (42) |
| IFI27 | Viral defense AM | Activated in viral infection via TLR7 activation | Tang et al., 2017 (43) |
| CTNNB1 | Intracellular bacterial defense AM | β-catenin-mediated bacterial elimination mechanism | Fu et al., 2017 (44) |
| SCD | Scavenger AM | There is high similarity between this cluster top DGEs and previously reported FABP4 expressing macrophages. | Coulombe et al., 2014 (46) |
| CTSC | M1-like, recruited, inflammatory AM |  |  |
| MSMO1 | Sterol synthesizing AM | Lipid metabolism gene signature | Poczobutt et al., 2016 (45) |
| SOD2 | Anti-cell-mediated inflammation AM |  |  |
| CD32 | Resident Ig-regulated AM |  |  |
| EIF | EIF+ AM | mTORC1 signaling pathway | Pugliese et al., 2017 (48) |
| G2/M Phase | G2/M-phase AM | Cycling macrophages | Travaglini et al., 2020 (49); Zilionis et al., 2019 (21) |
| S phase | S-phase AM | — |  |
| ISGs | Inflammatory AM | Similar gene expression observed in AM of severe COVID-19 patients | Liao et al., 2020 (29) |
| MTs | Metallothionein-expressing AM | Similar gene expression observed in AM of severe COVID-19 patients | Liao et al., 2020 (29) |

To further evaluate similarities and differences between AM subsets based on their ontogeny, we performed pseudotime trajectory analysis using Slingshot (37) (Fig 1D). This suggested that CD32b^+^ AMs have recently differentiated, whereas cycling AMs were found at the end of the trajectory. Next, we examined whether any of the identified AM clusters could be classified as M1- or M2-polarized macrophages based on known genes (38). In stable BAL samples, cluster 5 differentially expressed 4 genes associated with M1 polarization while cluster 8 differentially expressed 3 genes associated with M2 polarization (Fig S5C). Overall, however, expression of canonical M1- and M2-associated genes was distributed over multiple cell clusters, rather than being restricted to specific clusters. Since fetal yolk sac-derived and recruited monocyte-derived macrophages can contribute to the AM compartment, we further examined differential expression of genes associated with these populations (15). Interestingly, the M1-like CTSC+ cluster differentially expressed *CD14, CD74, APOE* and *C1QC* – which are associated with monocyte-derived macrophage populations. The M2-like CD32b+ cluster, by contrast, most strongly expressed *S100A10, CRIP1, S100A4*, and *MRC1* which are associated with tissue-resident yolk sac-derived AM (Fig 1E). Taken together, these observations suggest that the CTSC+ and CD32b+ clusters express genes classically associated with M1-like monocyte-derived AMs and M2-like resident AMs, respectively; however, these two populations are only a minority of AMs in LT BAL.

### Unique macrophages are found in BAL samples from patients with ALAD

Next we compared the transcriptomes of AMs from stable LT patients to those from patients with ALAD. As with the stable samples, we analyzed 1000 randomly selected AMs from ALAD samples and observed a similar degree of AM heterogeneity (Fig 2A, S7), with 13 distinct clusters. However, review of the top differentially expressed genes and pathway analysis (Fig S8) demonstrated that these clusters differed compared to those in stable samples. To identify similarities and differences between AM subsets in the two groups, we used Clustermap to compare differentially expressed genes by cluster from stable (data file S1) and ALAD (data file S2) samples. The two groups shared 11 common AM subsets, whereas there were two unique clusters in ALAD samples (Fig 2B-C and Fig S9A); these clusters expressed either interferon-stimulated genes (ISGs) or metallothioneins (MTs). Comparing the frequency of equivalent pairs of clusters in stable and ALAD samples, there were three additional clusters present at lower relative frequency (>20% difference in representation between the two groups) in stable samples (Fig 2C). In addition, while some ALAD AMs were in G2/M phase (Fig S9B), cells in this phase of the cell cycle only formed a distinct cluster in samples from stable patients. This observation suggests that there may be less transcriptional diversity among AMs in stable compared to ALAD samples, allowing greater resolution of S and G2/M phase AMs; alternatively, ALAD AMs might be more rapidly cycling, decreasing our ability to resolve differences between these phases of the cell cycle.

**Fig 2.**
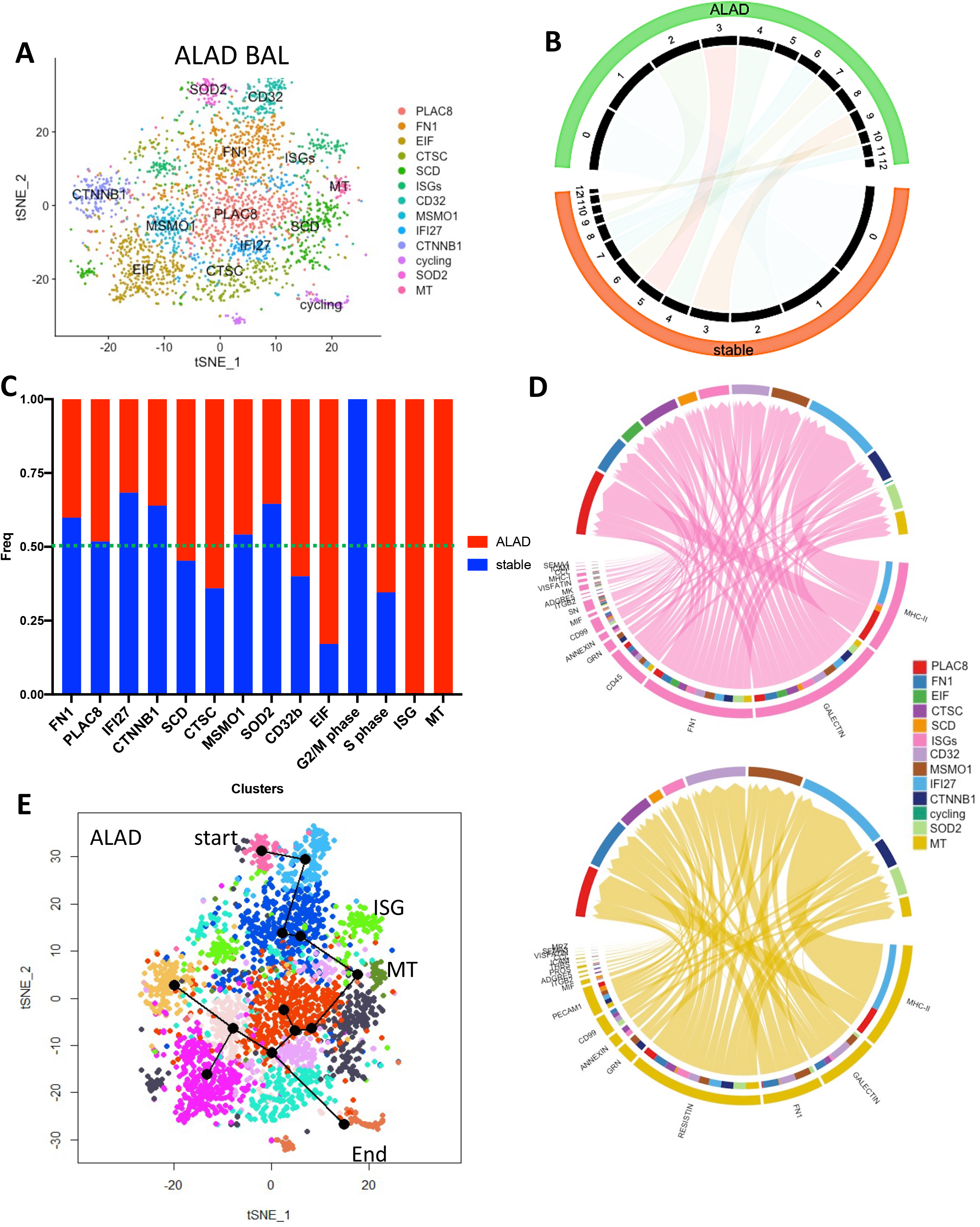
AMs with unique features are present in BAL from patients with acute lung allograft dysfunction. **(A)** tSNE plot of AMs from ALAD patients showing 13 distinct AM populations. Functional annotation of AMs was performed as in Fig 1B. **(B)** Clustermap analysis shows 11 AM clusters shared between ALAD (top) and stable (bottom) BAL. ALAD clusters 5 and 12 were unique to ALAD while stable cluster 10 was unique to stable samples. **(C)** Histogram showing representation of AM clusters in ALAD (red) and stable (blue) samples. Only ALAD samples contained ISG and MT AMs while only stable samples contained AMs forming a distinct G2/M-phase cluster. **(D)** Predicted cell-cell interaction networks of ISG (top) and MT (bottom) AMs with other AMs in ALAD BAL. Proteins involved in these interactions are listed along the bottom half of each semicircle, with the width of each band representing the proportion of cells in the ISG and MT clusters expressing each of the indicated molecules. The thickness of the bands joining proteins to other AM populations is proportional to the uniqueness of the predicted interactions. **(E)** Pseudotime trajectory analysis was performed using Slingshot. The pseudotime trajectories for stable and ALAD AMs were similar, with ALAD shown here. The start and end points of pseudotime are indicated, as are the positions of the ISG and MT clusters along the trajectory.

We delved deeper into the niche occupied by ISG and MT AMs by examining top differentially expressed genes, pathways and potential interactions of these cell clusters. Several highly expressed transcripts in the ISG cluster are involved in pro-inflammatory pathways, including *IFITM2, IFITM3, ISG15*, and *ISG20* in addition to *CXCL10*; the products of these genes promote lymphocyte trafficking and B cell activation in the transplantation setting (39) as well as in other inflammatory diseases (40). Our analysis showed that ISG AMs potentially interact with other AMs via a GALECTIN9-HAVCR2 pathway (Fig 2D top panel), which is known to promote macrophage-mediated inflammation (41, 42). Similarly, MT AMs are likely to interact with other AMs through the RESISTIN-CAP1 pathway, which induces a pro-inflammatory response in macrophages (43) (Fig 2D, bottom panel).

Next, we examined the differentiation of MT and ISG AMs using pseudotime analysis. This revealed that these cells arise at a similar stage (Fig 2E). Despite small differences in pseudotime branching, the general trajectory patterns for ALAD and stable samples were similar. In ALAD, MT and ISG AMs were in the early-intermediate part of the pseudotime trajectory, suggesting that they may have recently differentiated from recruited monocytes in response to specific stimuli.

### CXCL10+ AMs exhibit a pro-inflammatory phenotype *in vitro*

To determine whether AMs associated with ALAD could be identified based on cell-associated proteins, we obtained an independent set of 7 BAL samples (n=4 stable and n=3 ALAD) and stimulated the cells overnight in the presence or absence of lipopolysaccharide (LPS). We used flow cytometry to identify ISG AMs (full gating strategy is presented in Fig S10) by staining for CD163 and intracellular CXCL10. The percentage of CXCL10-expressing AMs was lower in BAL samples from stable patients but, with LPS stimulation, the proportion of CXCL10+ AMs in increased to the level seen in ALAD samples (Fig 3A). However, no further increase in CXCL10+ AMs was seen in ALAD samples in response to LPS, suggesting that only a limited number of AMs can acquire this phenotype. We analyzed cytokine and chemokine levels in the AM culture supernatants using a 13-parameter multiplexed cytokine assay. In keeping with the greater presence of CXCL10+ AMs in ALAD samples, IL-6, TNFα, IFNγ and CXCL10 were released in greater quantities by AMs from ALAD compared to stable samples (Fig 3B). These findings validate the observation that ISG AMs are associated with ALAD in LT patients.

**Fig 3.**
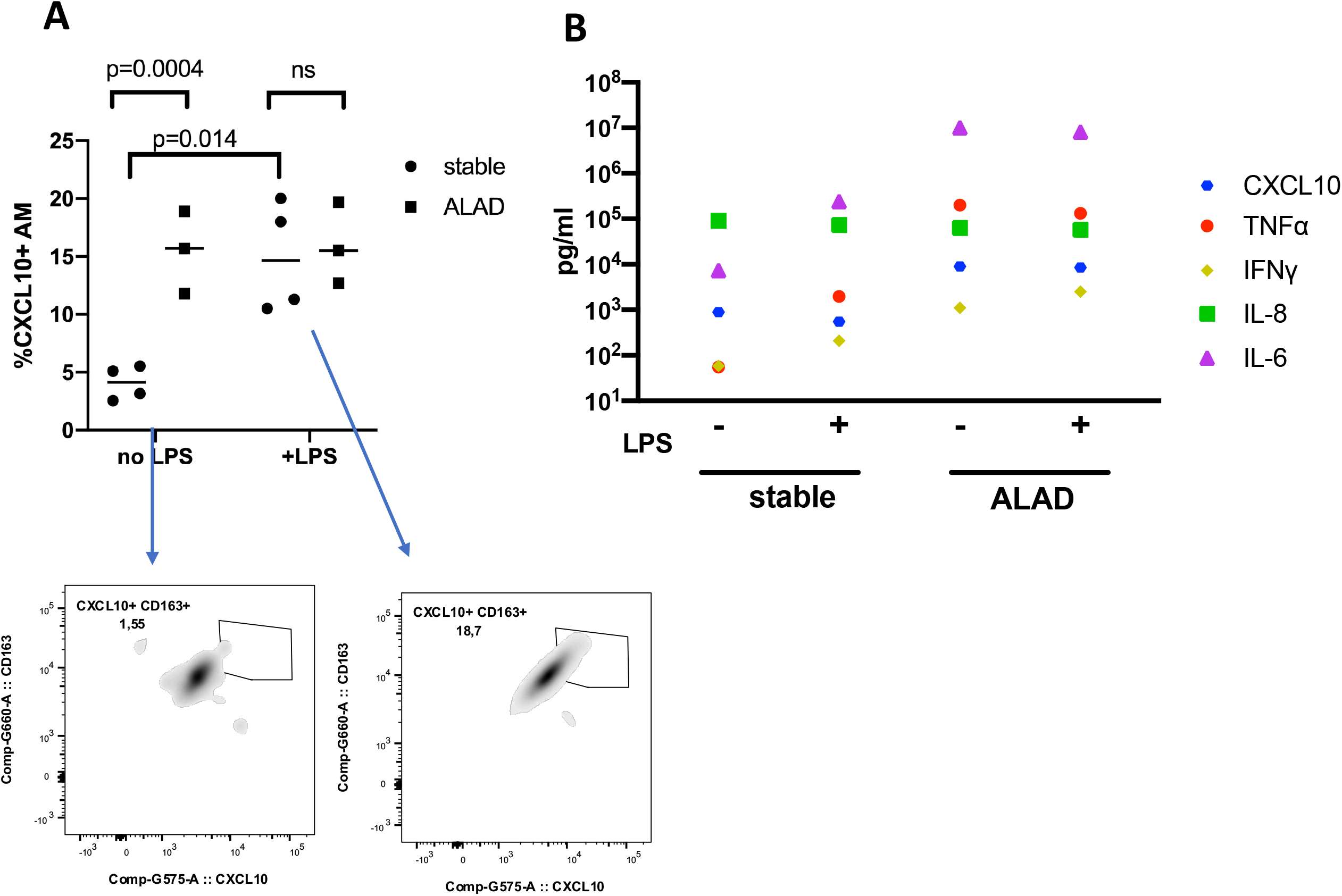
Identification of CXCL10-expressing AMs in an independent set of ALAD patients. In an independent set of BAL samples (n=4 stable and n=3 ALAD), AMs were cultured overnight in the presence or absence of LPS. AMs were identified as CD68+HLA-DR+ cells. **(A)** By intracellular staining, CXCL10 expression was elevated in AMs from ALAD compared to stable BAL samples; CXCL10 expression was augmented in stable AMs but was not further increased in ALAD AMs. Panels below show representative examples of CXCL10+CD163+ stable AMs without (left) and with (right) LPS stimulation. Repeated measures mixed effects model with Sidak’s post-hoc tests. **(B)**. Measurement of cytokine released into culture supernatants by freshly isolated BAL cells from 4 stable and 3 ALAD patients, with and without LPS stimulation. From 12 measured cytokines, IL-6, TNFa, IFNg and CXCL10 were released in greater quantities by AM from ALAD compared to stable samples. Data show the median of cytokine measurements for each group (n=4 stable and n=3 ALAD). Each sample was run in duplicate.

### Similarities and differences between AM subsets in ALAD and publicly available BAL datasets

To determine whether ISG and MT AMs are associated with inflammation outside the LT context, we investigated publicly available scRNAseq data from two studies, one focusing on the cellular composition of BAL from healthy controls (15) and another on BAL from COVID-19 patients (26). In healthy controls, Mould et al. described two macrophage clusters (m5 and m6) with features of ISG and MT AMs, respectively (15). We therefore integrated their data with our ALAD samples (Fig 4A). Based on expression level and cell frequency, the ALAD samples contributed more AMs to these clusters than did healthy control BAL samples (Fig 4B-C). We also found ISG and MT AMs in BAL samples from COVID-19 patients (clusters 0 and 1 for ISG and 22 for MT in Liao et al. (26)). In keeping with the notion that these AM populations mediate lung inflammation, most AMs in these clusters originated from patients with severe, rather than moderate, COVID-19 (Fig 4D-F). Clusters expressing the top differentially expressed genes in ISG and MT AMs are shown in Fig 4E-F, respectively.

**Fig 4.**
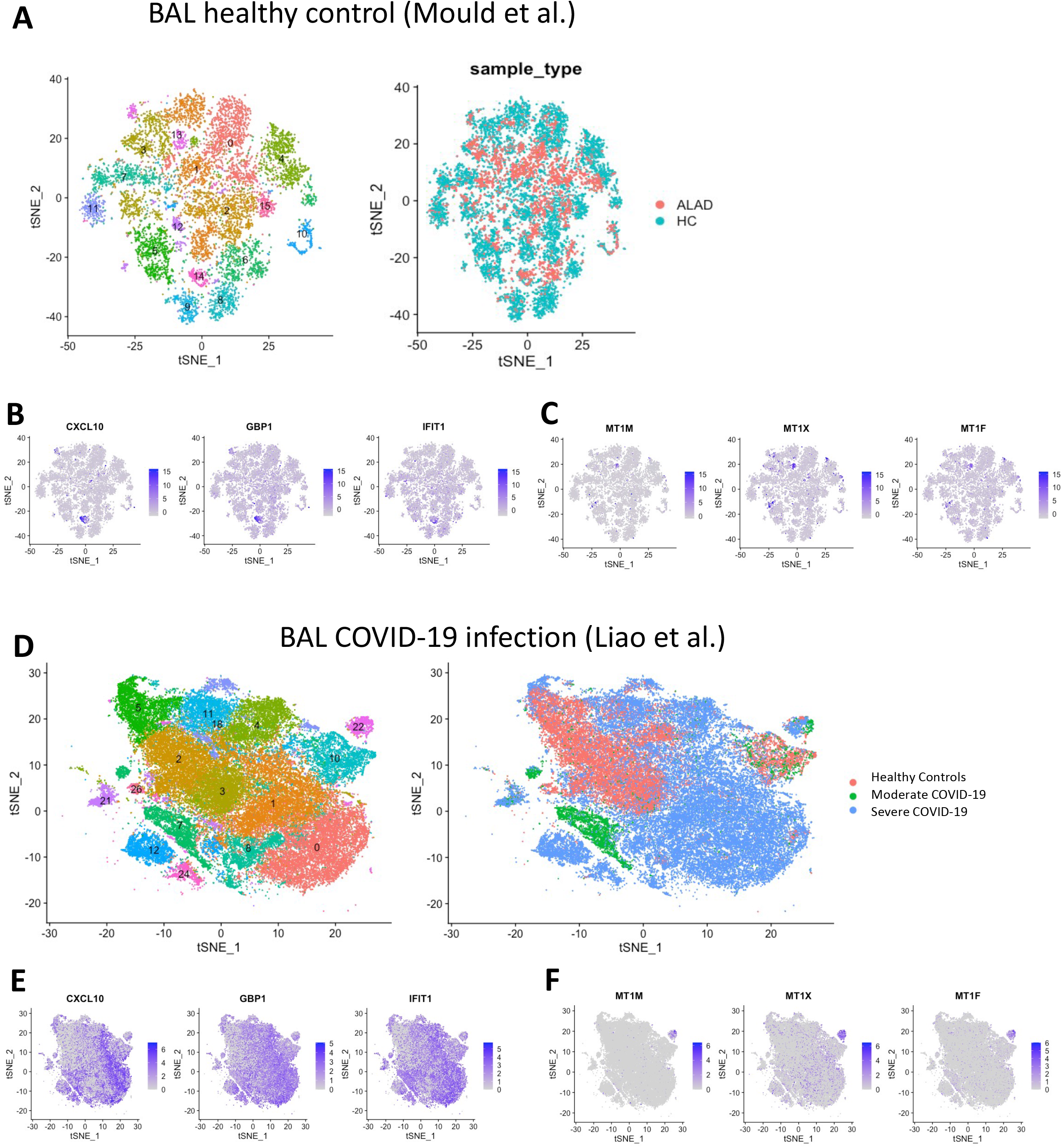
Association of ISG and MT AMs with inflammation in publicly available AM scRNAseq datasets. **(A)** tSNE plot of BAL AMs from 10 healthy controls (publicly available data from Mould et al., 2019) integrated with ALAD, grouped into distinct cell clusters (left panel) and grouped based on source of cells (right panel). **(B)** tSNE plot of integrated data shown in A highlighting top three genes differentially expressed by the ISG population (cluster 14 in panel A). ALAD BAL contributed more cells to this population than healthy controls (compare with A, right panel). **(C)** tSNE plot of integrated data shown in A highlighting top three genes differentially expressed by the MT population (cluster 13 in A). ALAD BAL contributed more cells to this population than healthy controls (compare with A, right panel). **(D)** tSNE plot of BAL AMs from COVID19 patients and healthy controls (publicly available data from Liao et al., 2020) grouped into distinct cell clusters (left panel) and grouped based on source of cells (right panel). **(E)** tSNE plot of data shown in D highlighting top three differentially expressed genes of the ISG clusters 0 and 1. Nearly all cells with this transcriptional profile originated from severe COVID-19 BAL rather than moderate COVID-19 or healthy control BAL. **(F)** tSNE plot from data shown in D highlighting the top three differentially expressed genes of the MT cluster 22, which also originated mostly from severe COVID-19 BAL.

### ALAD-associated macrophages are found in CLAD lung tissue

We next studied the transcriptomes of macrophages from fresh explanted CLAD lung tissue samples obtained at the time of re-transplantation for bronchiolitis obliterans syndrome (n=4) using the same approach taken stable BAL samples. CLAD tissue macrophages exhibited heterogeneity (Fig 5A). A review of the top differentially expressed genes (Fig S11) demonstrated that many of these clusters differed from BAL AMs, presumably because these samples contain both AMs and interstitial macrophages. Importantly, however, CLAD lung macrophages included distinct clusters with ISG and MT gene signatures (Fig 5B). The current consensus on healthy and diseased lung tissue macrophages holds that there are three main populations: FABP4^hi^, SPP1^hi^, and FCN1^hi^ (19, 44-46). Although these three genes are represented in different CLAD lung macrophages, they do not account for all BAL AMs and CLAD tissue macrophages in our samples or those of others (Fig 5C). Nevertheless, FABP4 expression – which has been associated with inflammation (44) – was clearly higher in ALAD BAL compared to stable samples and healthy control BAL from Mould et al. (15) (Fig 5C). We also observed that expression of SPP1 and FCN1 was greatest in macrophages from ALAD BAL and CLAD tissue, and nearly absent in BAL from stable LT recipients and healthy controls. While FABP4 seems to be expressed broadly across most ALAD and CLAD macrophages (Fig 5C, right column), the expression of SPP1 and FCN1 is more restricted and distinct from ISG and MT macrophages (c.f. Figs 1B, 2A and 5C, bottom row). Of all sample types, FCN1 and SPP1 were most prominent in CLAD lung tissue (Fig 5C, bottom row), suggesting that they may be primarily expressed in interstitial macrophages.

**Fig 5.**
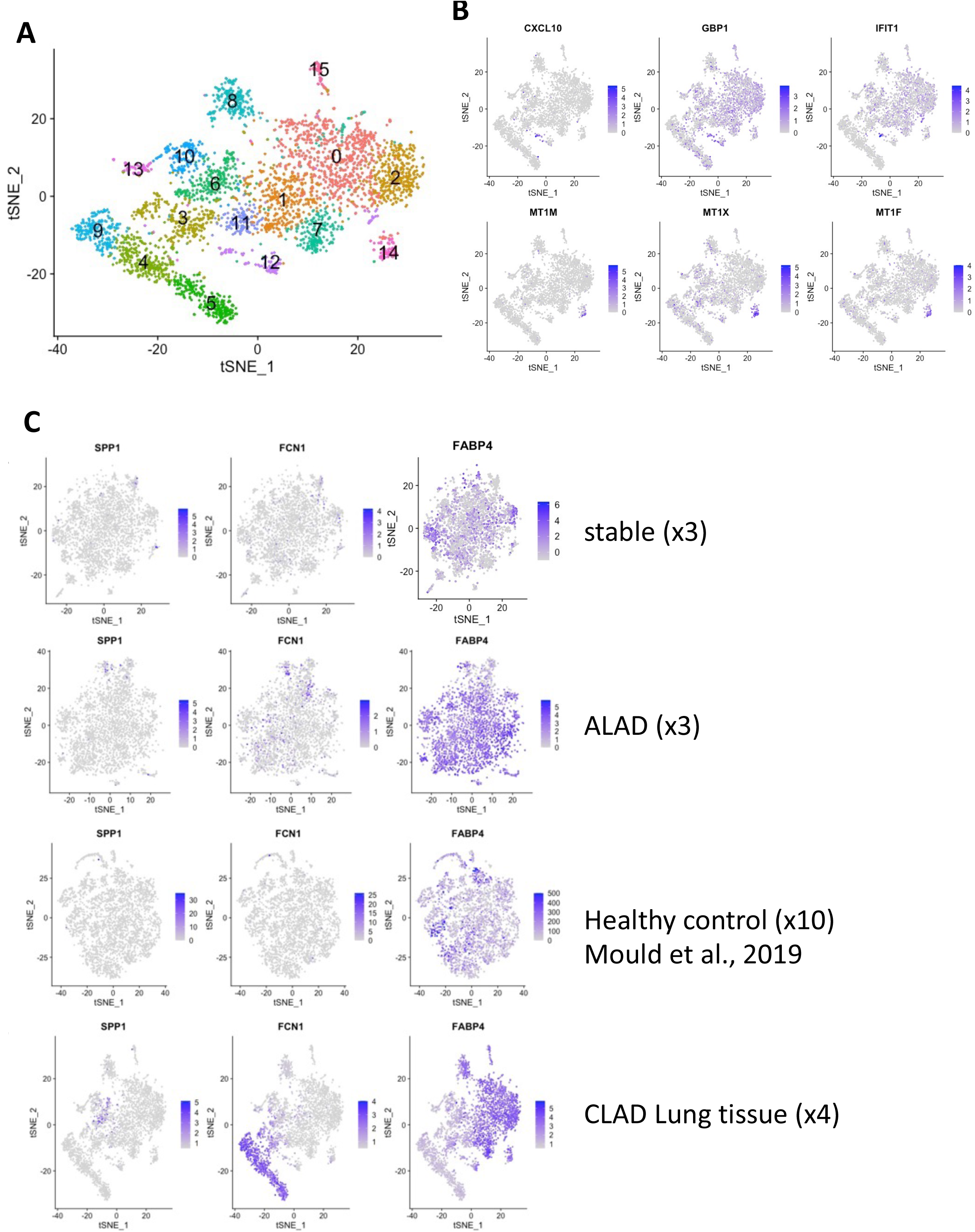
ISG and MT macrophages are present in CLAD lung tissue. **(A)** tSNE plot of macrophages from four integrated explanted CLAD lungs grouped into 16 distinct cell clusters. tSNE plots of integrated CLAD macrophages (shown in A) illustrating expression of ISG AM genes CXCL10, GBP1 and IFIT1 in CLAD macrophage cluster 12 (top row). A similar approach identified cluster 14 as MT macrophages in CLAD lung tissue (bottom row). **(C)** tSNE plots highlighting *SPP1, FCN1*, and *FABP4* gene expression in macrophages from stable (top row), ALAD (second row), healthy controls (from Mould et al. [ref 15], third row), and CLAD lung tissue (fourth row).

### Donor and recipient-derived cells contribute to all AM populations

Work by others (11, 36, 47, 48) and the data presented here illustrate that the AM compartment contains cycling cells that presumably permit self-renewal. In LT, this raises the question of whether and how long donor AMs persist in the allograft recipient. This issue has been addressed using HLA mismatching or detection of Y chromosomes in sex-mismatched LT, which have suggested an exceptionally long-term persistence of donor AM (49, 50). We chose to examine this question using an SNV calling algorithm that allowed us to infer donor and recipient origins of cells – without reference to genomic sequence data – by determining which SNVs were present in the majority of epithelial cells in the sample (Fig 6A). We observed replacement of donor-derived by recipient-derived AM over time post-transplant (Fig 6B), but in contrast to prior reports, most of the replacement occurs within the first-year post-transplant. At 3 months post-transplant, most AM were of donor origin; at 12 months post-transplant, only 10% of AM were donor-derived and by 24 months, AM were exclusively recipient-derived (Fig 6C). These findings reveal a somewhat faster replacement of donor AMs than has been reported previously.

**Fig 6.**
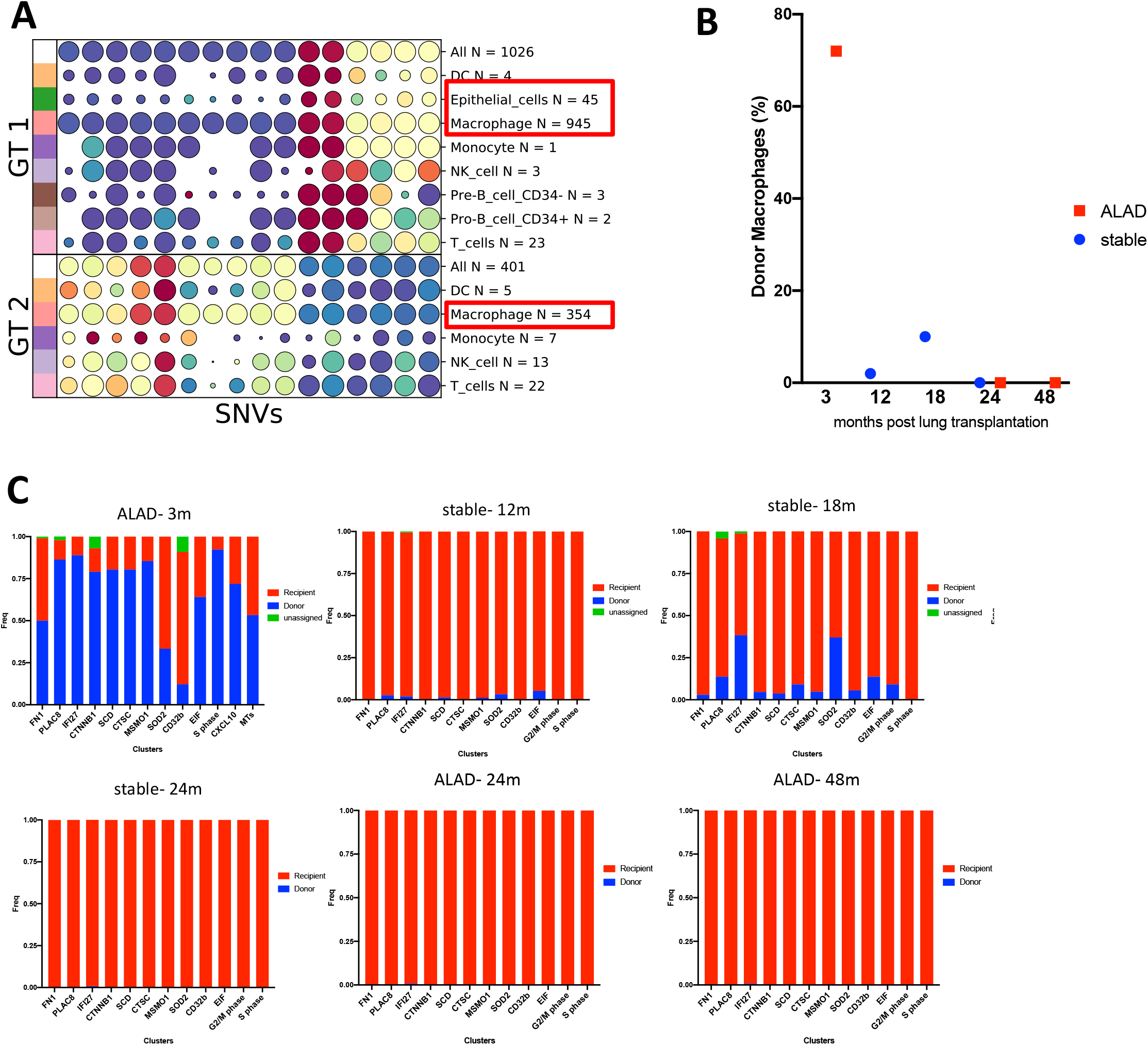
Identification of donor and recipient AMs using an SNV calling algorithm. **(A)** Identification of different SNVs within specific cell populations allowed identification of genotypes 1 and 2 (GT1 and GT2). In this example from an ALAD patient at 3 months post-LT, epithelial cells came from GT1 and are therefore ascribed to the donor. AMs of GT1 outnumbered those of GT2 (red boxes). Circle sizes indicate the proportion of cells in which each SNV is expressed, whereas colours indicate the average minor allele fraction across the cells in each group (red=high, blue=low). **(B)** In both ALAD (red) and stable (blue) BAL samples, the proportion of donor AMs decreased with time post-LT. **(C)** Histograms showing the proportions of donor (blue) and recipient (red) AMs in BAL samples at various times post-transplant.

## Discussion

In this study, we used scRNAseq to compare macrophage transcriptional states in BAL samples from LT recipients with either stable lung function or ALAD, and from explanted CLAD lung tissue. We focused on macrophages because they represent the majority of BAL cells, and the behaviour of AMs in relation to lung allograft dysfunction has not been investigated in detail. Our data reveal a previously unappreciated diversity in LT AMs, with evidence for functional specialization among the populations. In agreement with previous reports (18, 51, 52), our data show that the M1 vs. M2 paradigm inadequately captures AM diversity.

We observed substantial differences – despite our small sample size and the high genetic diversity associated with human subjects – between stable and ALAD patients. Most notably, ISG and MT AMs were uniquely represented in ALAD samples. An examination of differentially expressed genes in ISG AMs suggested that different pro-inflammatory cascades have been induced, probably in response to IFNγ, including guanylate-binding proteins (GBPs), cytokines and chemokines (CXCL10, CCL4, and CCL2), in addition to several members of the ISG family. Although the IFNγ response in macrophages is typically associated with infection, it is likely that ISG AMs in our ALAD samples were responding to alloimmune-mediated inflammation since patients with infection were excluded. However, it remains unclear whether this cluster of cells represents a developmentally distinct macrophage population, or whether the activation of an ISG program simply reflects the effects of IFNγ on a susceptible subset of AMs. Since these cells were absent from stable BAL samples that underwent scRNAseq and only represented in small proportions of stable samples subjected to flow cytometry, we favour the latter explanation. Importantly, AMs isolated from stable patients produced less CXCL10 – one of the most differentially expressed genes in the ISG cluster – than AMs from ALAD patients. This finding validates the scRNAseq observations at a phenotypic level. Nevertheless, it was possible to elicit CXCL10 expression in stable AMs that was comparable to that of ALAD AMs using LPS, indicating that the AM phenotypes we describe here are dynamic.

MT AMs – also uniquely associated with ALAD – have not previously been shown to participate in alloimmunity. MTs are a family of metal-binding proteins that maintain homeostasis of zinc and copper, mitigate heavy metal toxication, and alleviate superoxide stress. They also mediate inflammatory responses and antimicrobial defense (53). It has been suggested that MT expression in macrophages is required for pro-inflammatory responses (54). We are unaware of previous data implicating MT-expressing macrophages in LT; whether they participate in the pathogenesis of allograft dysfunction requires further investigation.

Nevertheless, our observation that ISG and MT macrophages are present in CLAD lung tissue confirms their relationship to progressive allograft dysfunction. Further, that these cellular populations are associated with disease severity in COVID-19 and are present in only small numbers in healthy lungs supports their role in lung inflammation more generally. Identification of specific disease-associated macrophages may facilitate the discovery of targeted therapies for inflammatory lung diseases and lung allograft dysfunction in the future.

In support of this notion, we also observed remarkably extensive cell-cell communication networks amongst AMs in both stable and ALAD patients, suggesting the existence of complex feedback and feed-forward loops between these cells in the bronchoalveolar niche; in contrast, communication between AMs and T cells appeared to be more limited, with predicted pathways primarily restricted to chemokine signalling. These findings will require further study, but they suggest that control of the inflammatory tone of the alveolar space in LT recipients is subject primarily to complex inter-relationships between AMs rather than AM-T cell interactions.

Our SNV data confirmed previous reports (49) describing replacement of donor by recipient AMs following LT. While prior investigations suggested that donor AM replacement takes years, we found that donor AMs are mostly replaced within 12 months post-LT. The previous work distinguished donor and recipient AMs using flow cytometric detection of allogeneic HLA molecules, whereas we used SNVs. We believe that the latter, with its coverage of the sequenced transcriptome, more accurately attributes each cell to its origin than HLA typing, since intact allogeneic HLA molecules can move from one cell to another via trogocytosis or exosome-mediated transfer (55). Our findings require further validation, but suggest that relying on surface expression of allogeneic HLA molecules might overestimate donor AM persistence. This is an important area for future investigation, since the rate of clearance of donor leukocytes over time may have prognostic implications in LT (56).

In summary, we have shown that the AM compartment undergoes specific transcriptional alterations during ALAD, with primarily recipient-origin monocyte-derived ISG and MT AMs emerging. To date, most research on ALAD has focused on lymphocyte-mediated adaptive immune responses. While these are undoubtedly central to lung allograft loss, the appearance of specific AM gene expression programs during ALAD suggests that these cells may contribute actively to the loss of lung function and may therefore represent important mechanistic targets.

## Acknowledgements

The authors wish to thank Marcelo Cuesta, Iva Avramov and Max Niit of the Toronto Lung Transplant Program Biobank for assistance in sample handling. We thank Amber Xue for help with the multiplexed cytokine assay. We also thank Gurbaksh Basi and Iulia Cirlan of the Princess Margaret Genomics Centre for running scRNAseq.

## Funding

This work was supported by a Cystic Fibrosis Foundation Mechanisms of Chronic Lung Allograft Dysfunction grant (#JUVET18AB0) to S.J., a Sanofi iAward to S.J. and T.M., and a Canadian Society of Transplantation Fellowship (to S.M.).

## Author contributions

**S.M**. conceived of and performed experiments, analyzed data and wrote the manuscript. **A.D**. performed experiments and analyzed data.**G.W**. analyzed data and wrote the manuscript. **A.D**. performed experiments and analyzed data. **T.A**. analyzed data. **G.B**. collected and analyzed data. **B.R-P**. collected and analyzed data. **S.K**. conceived of experiments and wrote the manuscript. **S.MacP**. analyzed data and wrote the manuscript. **M.L**. contributed to analysis and wrote the manuscript. **J.Y**. analyzed data and wrote the manuscript. **T.M**. contributed to data analysis and wrote the manuscript. **S.J**. conceived of experiments, analyzed data and wrote the manuscript.

## Competing interests

none declared.

## Data availability

Data and codes are available in https://github.com/SamWell16/BAL-scRNAseq.

## Notes

### Competing Interest Statement

The authors have declared no competing interest.

## References

1. Hachem RR. The role of the immune system in lung transplantation: towards improved long-term results. J Thorac Dis 2019; 11: S1721–S1731.

2. Martinu T, Koutsokera A, Benden C, Cantu E, Chambers D, Cypel M, Edelman J, Emtiazjoo A, Fisher AJ, Greenland JR, Hayes D, Jr., Hwang D, Keller BC, Lease ED, Perch M, Sato M, Todd JL, Verleden S, von der Thusen J, Weigt SS, Keshavjee S, bronchoalveolar lavage standardization w. International Society for Heart and Lung Transplantation consensus statement for the standardization of bronchoalveolar lavage in lung transplantation. J Heart Lung Transplant 2020.

3. Kennedy VE, Todd JL, Palmer SM. Bronchoalveolar lavage as a tool to predict, diagnose and understand bronchiolitis obliterans syndrome. Am J Transplant 2013; 13: 552–561.

4. Speck NE, Schuurmans MM, Murer C, Benden C, Huber LC. Diagnostic value of plasma and bronchoalveolar lavage samples in acute lung allograft rejection: differential cytology. Respir Res 2016; 17: 74.

5. Vos R, Vanaudenaerde BM, Verleden SE, De Vleeschauwer SI, Willems-Widyastuti A, Van Raemdonck DE, Dupont LJ, Nawrot TS, Verbeken EK, Verleden GM. Bronchoalveolar lavage neutrophilia in acute lung allograft rejection and lymphocytic bronchiolitis. J Heart Lung Transplant 2010; 29: 1259–1269.

6. Greenland JR, Jewell NP, Gottschall M, Trivedi NN, Kukreja J, Hays SR, Singer JP, Golden JA, Caughey GH. Bronchoalveolar lavage cell immunophenotyping facilitates diagnosis of lung allograft rejection. Am J Transplant 2014; 14: 831–840.

7. Herold S, Mayer K, Lohmeyer J. Acute lung injury: how macrophages orchestrate resolution of inflammation and tissue repair. Front Immunol 2011; 2: 65.

8. Westphalen K, Gusarova GA, Islam MN, Subramanian M, Cohen TS, Prince AS, Bhattacharya J. Sessile alveolar macrophages communicate with alveolar epithelium to modulate immunity. Nature 2014; 506: 503–506.

9. Hussell T, Bell TJ. Alveolar macrophages: plasticity in a tissue-specific context. Nat Rev Immunol 2014; 14: 81–93.

10. Byrne AJ, Mathie SA, Gregory LG, Lloyd CM. Pulmonary macrophages: key players in the innate defence of the airways. Thorax 2015; 70: 1189–1196.

11. Hashimoto D, Chow A, Noizat C, Teo P, Beasley MB, Leboeuf M, Becker CD, See P, Price J, Lucas D, Greter M, Mortha A, Boyer SW, Forsberg EC, Tanaka M, van Rooijen N, Garcia-Sastre A, Stanley ER, Ginhoux F, Frenette PS, Merad M. Tissue-resident macrophages self-maintain locally throughout adult life with minimal contribution from circulating monocytes. Immunity 2013; 38: 792–804.

12. van de Laar L, Saelens W, De Prijck S, Martens L, Scott CL, Van Isterdael G, Hoffmann E, Beyaert R, Saeys Y, Lambrecht BN, Guilliams M. Yolk Sac Macrophages, Fetal Liver, and Adult Monocytes Can Colonize an Empty Niche and Develop into Functional Tissue-Resident Macrophages. Immunity 2016; 44: 755–768.

13. Svedberg FR, Brown SL, Krauss MZ, Campbell L, Sharpe C, Clausen M, Howell GJ, Clark H, Madsen J, Evans CM, Sutherland TE, Ivens AC, Thornton DJ, Grencis RK, Hussell T, Cunoosamy DM, Cook PC, MacDonald AS. The lung environment controls alveolar macrophage metabolism and responsiveness in type 2 inflammation. Nat Immunol 2019; 20: 571–580.

14. Byrne AJ, Powell JE, O’Sullivan BJ, Ogger PP, Hoffland A, Cook J, Bonner KL, Hewitt RJ, Wolf S, Ghai P, Walker SA, Lukowski SW, Molyneaux PL, Saglani S, Chambers DC, Maher TM, Lloyd CM. Dynamics of human monocytes and airway macrophages during healthy aging and after transplant. J Exp Med 2020; 217.

15. Mould KJ, Jackson ND, Henson PM, Seibold M, Janssen WJ. Single cell RNA sequencing identifies unique inflammatory airspace macrophage subsets. JCI Insight 2019; 4.

16. Peyser R, MacDonnell S, Gao Y, Cheng L, Kim Y, Kaplan T, Ruan Q, Wei Y, Ni M, Adler C, Zhang W, Devalaraja-Narashimha K, Grindley J, Halasz G, Morton L. Defining the Activated Fibroblast Population in Lung Fibrosis Using Single-Cell Sequencing. Am J Respir Cell Mol Biol 2019; 61: 74–85.

17. Lukassen S, Chua RL, Trefzer T, Kahn NC, Schneider MA, Muley T, Winter H, Meister M, Veith C, Boots AW, Hennig BP, Kreuter M, Conrad C, Eils R. SARS-CoV-2 receptor ACE2 and TMPRSS2 are primarily expressed in bronchial transient secretory cells. EMBO J 2020; 39: e105114.

18. Zilionis R, Engblom C, Pfirschke C, Savova V, Zemmour D, Saatcioglu HD, Krishnan I, Maroni G, Meyerovitz CV, Kerwin CM, Choi S, Richards WG, De Rienzo A, Tenen DG, Bueno R, Levantini E, Pittet MJ, Klein AM. Single-Cell Transcriptomics of Human and Mouse Lung Cancers Reveals Conserved Myeloid Populations across Individuals and Species. Immunity 2019; 50: 1317–1334 e1310.

19. Reyfman PA, Walter JM, Joshi N, Anekalla KR, McQuattie-Pimentel AC, Chiu S, Fernandez R, Akbarpour M, Chen CI, Ren Z, Verma R, Abdala-Valencia H, Nam K, Chi M, Han S, Gonzalez-Gonzalez FJ, Soberanes S, Watanabe S, Williams KJN, Flozak AS, Nicholson TT, Morgan VK, Winter DR, Hinchcliff M, Hrusch CL, Guzy RD, Bonham CA, Sperling AI, Bag R, Hamanaka RB, Mutlu GM, Yeldandi AV, Marshall SA, Shilatifard A, Amaral LAN, Perlman H, Sznajder JI, Argento AC, Gillespie CT, Dematte J, Jain M, Singer BD, Ridge KM, Lam AP, Bharat A, Bhorade SM, Gottardi CJ, Budinger GRS, Misharin AV. Single-Cell Transcriptomic Analysis of Human Lung Provides Insights into the Pathobiology of Pulmonary Fibrosis. Am J Respir Crit Care Med 2019; 199: 1517–1536.

20. Schiller HB, Montoro DT, Simon LM, Rawlins EL, Meyer KB, Strunz M, Vieira Braga FA, Timens W, Koppelman GH, Budinger GRS, Burgess JK, Waghray A, van den Berge M, Theis FJ, Regev A, Kaminski N, Rajagopal J, Teichmann SA, Misharin AV, Nawijn MC. The Human Lung Cell Atlas: A High-Resolution Reference Map of the Human Lung in Health and Disease. Am J Respir Cell Mol Biol 2019; 61: 31–41.

21. Zou X, Chen K, Zou J, Han P, Hao J, Han Z. Single-cell RNA-seq data analysis on the receptor ACE2 expression reveals the potential risk of different human organs vulnerable to 2019-nCoV infection. Front Med 2020; 14: 185–192.

22. Kim N, Kim HK, Lee K, Hong Y, Cho JH, Choi JW, Lee JI, Suh YL, Ku BM, Eum HH, Choi S, Choi YL, Joung JG, Park WY, Jung HA, Sun JM, Lee SH, Ahn JS, Park K, Ahn MJ, Lee HO. Single-cell RNA sequencing demonstrates the molecular and cellular reprogramming of metastatic lung adenocarcinoma. Nat Commun 2020; 11: 2285.

23. Aran D, Looney AP, Liu L, Wu E, Fong V, Hsu A, Chak S, Naikawadi RP, Wolters PJ, Abate AR, Butte AJ, Bhattacharya M. Reference-based analysis of lung single-cell sequencing reveals a transitional profibrotic macrophage. Nat Immunol 2019; 20: 163–172.

24. Schyns J, Bai Q, Ruscitti C, Radermecker C, De Schepper S, Chakarov S, Farnir F, Pirottin D, Ginhoux F, Boeckxstaens G, Bureau F, Marichal T. Non-classical tissue monocytes and two functionally distinct populations of interstitial macrophages populate the mouse lung. Nat Commun 2019; 10: 3964.

25. Bost P, Giladi A, Liu Y, Bendjelal Y, Xu G, David E, Blecher-Gonen R, Cohen M, Medaglia C, Li H, Deczkowska A, Zhang S, Schwikowski B, Zhang Z, Amit I. Host-Viral Infection Maps Reveal Signatures of Severe COVID-19 Patients. Cell 2020; 181: 1475–1488 e1412.

26. Liao M, Liu Y, Yuan J, Wen Y, Xu G, Zhao J, Cheng L, Li J, Wang X, Wang F, Liu L, Amit I, Zhang S, Zhang Z. Single-cell landscape of bronchoalveolar immune cells in patients with COVID-19. Nat Med 2020; 26: 842–844.

27. Jin S, Guerrero-Juarez CF, Zhang L, Chang I, Myung P, Plikus MV, Nie Q. Inference and analysis of cell-cell communication using CellChat. bioRxiv 2020: 2020.2007.2021.214387.

28. Stuart T, Butler A, Hoffman P, Hafemeister C, Papalexi E, Mauck WM, 3rd, Hao Y, Stoeckius M, Smibert P, Satija R. Comprehensive Integration of Single-Cell Data. Cell 2019; 177: 1888–1902 e1821.

29. Ledford JG, Kovarova M, Koller BH. Impaired host defense in mice lacking ONZIN. J Immunol 2007; 178: 5132–5143.

30. Tang BM, Shojaei M, Parnell GP, Huang S, Nalos M, Teoh S, O’Connor K, Schibeci S, Phu AL, Kumar A, Ho J, Meyers AFA, Keynan Y, Ball T, Pisipati A, Kumar A, Moore E, Eisen D, Lai K, Gillett M, Geffers R, Luo H, Gul F, Schreiber J, Riedel S, Booth D, McLean A, Schughart K. A novel immune biomarker IFI27 discriminates between influenza and bacteria in patients with suspected respiratory infection. Eur Respir J 2017; 49.

31. Fu Q, Chen K, Zhu Q, Wang W, Huang F, Miao L, Wu X. beta-catenin promotes intracellular bacterial killing via suppression of Pseudomonas aeruginosa-triggered macrophage autophagy. J Int Med Res 2017; 45: 556–569.

32. Poczobutt JM, D. S, Yadav VK, Nguyen TT, Li H, Sippel TR, Weiser-Evans MC, Nemenoff RA. Expression Profiling of Macrophages Reveals Multiple Populations with Distinct Biological Roles in an Immunocompetent Orthotopic Model of Lung Cancer. J Immunol 2016; 196: 2847–2859.

33. Coulombe F, Jaworska J, Verway M, Tzelepis F, Massoud A, Gillard J, Wong G, Kobinger G, Xing Z, Couture C, Joubert P, Fritz JH, Powell WS, Divangahi M. Targeted prostaglandin E2 inhibition enhances antiviral immunity through induction of type I interferon and apoptosis in macrophages. Immunity 2014; 40: 554–568.

34. Jablonski KA, Amici SA, Webb LM, Ruiz-Rosado Jde D, Popovich PG, Partida-Sanchez S, Guerau-de-Arellano M. Novel Markers to Delineate Murine M1 and M2 Macrophages. PLoS One 2015; 10: e0145342.

35. Pugliese SC, Kumar S, Janssen WJ, Graham BB, Frid MG, Riddle SR, El Kasmi KC, Stenmark KR. A Time- and Compartment-Specific Activation of Lung Macrophages in Hypoxic Pulmonary Hypertension. J Immunol 2017; 198: 4802–4812.

36. Travaglini KJ, Nabhan AN, Penland L, Sinha R, Gillich A, Sit RV, Chang S, Conley SD, Mori Y, Seita J, Berry GJ, Shrager JB, Metzger RJ, Kuo CS, Neff N, Weissman IL, Quake SR, Krasnow MA. A molecular cell atlas of the human lung from single-cell RNA sequencing. Nature 2020; 587: 619–625.

37. Street K, Risso D, Fletcher RB, Das D, Ngai J, Yosef N, Purdom E, Dudoit S. Slingshot: cell lineage and pseudotime inference for single-cell transcriptomics. BMC Genomics 2018; 19: 477.

38. Orecchioni M, Ghosheh Y, Pramod AB, Ley K. Macrophage Polarization: Different Gene Signatures in M1(LPS+) vs. Classically and M2(LPS-) vs. Alternatively Activated Macrophages. Front Immunol 2019; 10: 1084.

39. Hakim FT, Memon S, Jin P, Imanguli MM, Wang H, Rehman N, Yan XY, Rose J, Mays JW, Dhamala S, Kapoor V, Telford W, Dickinson J, Davis S, Halverson D, Naik HB, Baird K, Fowler D, Stroncek D, Cowen EW, Pavletic SZ, Gress RE. Upregulation of IFN-Inducible and Damage-Response Pathways in Chronic Graft-versus-Host Disease. J Immunol 2016; 197: 3490–3503.

40. Zhang F, Mears JR, Shakib L, Beynor JI, Shanaj S, Korsunsky I, Nathan A, Accelerating Medicines Partnership Rheumatoid A, Systemic Lupus Erythematosus C, Donlin LT, Raychaudhuri S. IFN-gamma and TNF-alpha drive a CXCL10+ CCL2+ macrophage phenotype expanded in severe COVID-19 lungs and inflammatory diseases with tissue inflammation. Genome Med 2021; 13: 64.

41. Sada-Ovalle I, Chavez-Galan L, Torre-Bouscoulet L, Nava-Gamino L, Barrera L, Jayaraman P, Torres-Rojas M, Salazar-Lezama MA, Behar SM. The Tim3-galectin 9 pathway induces antibacterial activity in human macrophages infected with Mycobacterium tuberculosis. J Immunol 2012; 189: 5896–5902.

42. Ma CJ, Li GY, Cheng YQ, Wang JM, Ying RS, Shi L, Wu XY, Niki T, Hirashima M, Li CF, Moorman JP, Yao ZQ. Cis association of galectin-9 with Tim-3 differentially regulates IL-12/IL-23 expressions in monocytes via TLR signaling. PLoS One 2013; 8: e72488.

43. Silswal N, Singh AK, Aruna B, Mukhopadhyay S, Ghosh S, Ehtesham NZ. Human resistin stimulates the pro-inflammatory cytokines TNF-alpha and IL-12 in macrophages by NF-kappaB-dependent pathway. Biochem Biophys Res Commun 2005; 334: 1092–1101.

44. Morse C, Tabib T, Sembrat J, Buschur KL, Bittar HT, Valenzi E, Jiang Y, Kass DJ, Gibson K, Chen W, Mora A, Benos PV, Rojas M, Lafyatis R. Proliferating SPP1/MERTK-expressing macrophages in idiopathic pulmonary fibrosis. Eur Respir J 2019; 54.

45. Adams TS, Schupp JC, Poli S, Ayaub EA, Neumark N, Ahangari F, Chu SG, Raby BA, DeIuliis G, Januszyk M, Duan Q, Arnett HA, Siddiqui A, Washko GR, Homer R, Yan X, Rosas IO, Kaminski N. Single-cell RNA-seq reveals ectopic and aberrant lung-resident cell populations in idiopathic pulmonary fibrosis. Sci Adv 2020; 6: eaba1983.

46. Schupp JC, Khanal S, Gomez JL, Sauler M, Adams TS, Chupp GL, Yan X, Poli S, Zhao Y, Montgomery RR, Rosas IO, Dela Cruz CS, Bruscia EM, Egan ME, Kaminski N, Britto CJ. Single-Cell Transcriptional Archetypes of Airway Inflammation in Cystic Fibrosis. Am J Respir Crit Care Med 2020; 202: 1419–1429.

47. Ginhoux F, Greter M, Leboeuf M, Nandi S, See P, Gokhan S, Mehler MF, Conway SJ, Ng LG, Stanley ER, Samokhvalov IM, Merad M. Fate mapping analysis reveals that adult microglia derive from primitive macrophages. Science 2010; 330: 841–845.

48. Yona S, Kim KW, Wolf Y, Mildner A, Varol D, Breker M, Strauss-Ayali D, Viukov S, Guilliams M, Misharin A, Hume DA, Perlman H, Malissen B, Zelzer E, Jung S. Fate mapping reveals origins and dynamics of monocytes and tissue macrophages under homeostasis. Immunity 2013; 38: 79–91.

49. Nayak DK, Zhou F, Xu M, Huang J, Tsuji M, Hachem R, Mohanakumar T. Long-Term Persistence of Donor Alveolar Macrophages in Human Lung Transplant Recipients That Influences Donor-Specific Immune Responses. Am J Transplant 2016; 16: 2300–2311.

50. Eguiluz-Gracia I, Schultz HH, Sikkeland LI, Danilova E, Holm AM, Pronk CJ, Agace WW, Iversen M, Andersen C, Jahnsen FL, Baekkevold ES. Long-term persistence of human donor alveolar macrophages in lung transplant recipients. Thorax 2016; 71: 1006–1011.

51. Nahrendorf M, Swirski FK. Abandoning M1/M2 for a Network Model of Macrophage Function. Circ Res 2016; 119: 414–417.

52. Evren E, Ringqvist E, Willinger T. Origin and ontogeny of lung macrophages: from mice to humans. Immunology 2020; 160: 126–138.

53. Subramanian Vignesh K, Deepe GS, Jr. Metallothioneins: Emerging Modulators in Immunity and Infection. Int J Mol Sci 2017; 18.

54. Kanekiyo M, Itoh N, Kawasaki A, Matsuda K, Nakanishi T, Tanaka K. Metallothionein is required for zinc-induced expression of the macrophage colony stimulating factor gene. J Cell Biochem 2002; 86: 145–153.

55. Liu Q, Rojas-Canales DM, Divito SJ, Shufesky WJ, Stolz DB, Erdos G, Sullivan ML, Gibson GA, Watkins SC, Larregina AT, Morelli AE. Donor dendritic cell-derived exosomes promote allograft-targeting immune response. J Clin Invest 2016; 126: 2805–2820.

56. Snyder ME, Finlayson MO, Connors TJ, Dogra P, Senda T, Bush E, Carpenter D, Marboe C, Benvenuto L, Shah L, Robbins H, Hook JL, Sykes M, D’Ovidio F, Bacchetta M, Sonett JR, Lederer DJ, Arcasoy S, Sims PA, Farber DL. Generation and persistence of human tissue-resident memory T cells in lung transplantation. Sci Immunol 2019; 4.

