## Supplementary materials and figures for "Interferon-stimulated and metallothionein-expressing macrophages are associated with acute and chronic allograft dysfunction after lung transplantation"

### Supplementary materials and methods

Isolation of BAL cells: BAL samples were centrifuged at 100*g* for 5 minutes at 4°C to pellet larger and more fragile epithelial cells. The supernatant was then transferred to a new tube for a second centrifugation at 400*g* for 5 minutes at 4°C to pellet the remaining cells. The two cell pellets were combined by resuspending in 500 ul of cold PBS + 0.5% fetal bovine serum. Cells were counted and transported to the UHN genomics center for library preparation within 30 minutes of BAL sample acquisition.

Isolation of CLAD lung cells: CLAD lungs were collected immediately after explantation from patients undergoing re-transplantation and were flushed with 500mL saline via the pulmonary vein. After flush, a wedge (16cm x 8cm x 1cm) was cut from the upper lobe. From the center of this wedge, a 4cm x 4cm x 1cm piece of lung tissue was collected, minced and submerged into a digestion solution containing 0.80mg/ml of Collagenase A (Roche) and 0.16mg/ml DNase I (Roche) in HBSS (gibco) plus 5% v/v heat-inactivated fetal bovine serum (Wisent) and 10nM HEPES (Thermo) and dissociated with a gentleMACS™ Octo Dissociator (Miltenyi Biotec) for 10min at 37°C, followed by the "liver_4" setting. The dissociate was strained with a 70µm cell strainer, and red blood cells were lysed with ACK buffer. Sample were washed with PBS (Wisent) + 0.04% BSA (Sigma), and cell count and viability were assessed with hemocytometer.

Capture, library construction and sequencing: A total of 11 µl of the single cell suspension and 40 µl barcoded Gel Beads were loaded onto a Chromium Chip A (10x Genomics) to generate single-cell gel bead-in-emulsion. Single cell capture and downstream library construction were performed using 10x Genomics 3' expression V2 chemistry. Full-length cDNA along with cell-barcode identifiers were PCR-amplified and sequencing libraries were prepared. The constructed library was frozen for later sequencing on the Illumina® NovaSeq platform.

scRNAseq data alignment and quality control: Cell Ranger (27) was used to perform sample de-multiplexing, barcode processing and single-cell 3' UMI counting with human GRCh38 as the reference genome. Specifically, splicing-aware aligner STAR was used for FASTQ alignment. Cell barcodes were then determined based on distribution of UMI count automatically. Each individual sample was run through Seurat v3 (28) with the following filtration criteria applied to each cell: gene number between 200 and 5000, UMI count between 1000 and 30,000 and mitochondrial gene percentage below 10%. Prior to downstream analysis, we also removed genes for ribosomal proteins (those whose names started with RPL or RPS) as suggested by Mould and associates (15). After filtration, the remaining cells (stable, 3044 ± 1519 cells; ALAD, 2593 ± 904 cells) were subjected to further analysis.

Cell annotation: Cells were normalized using the "log normalization" method in Seurat v3 and run through the SingleR package (23) to annotate each individual cell in comparison to the Human Primary Cell Atlas (55). Cellular annotation was added to the Seurat object as meta-data.

In the Seurat object of each individual sample, cells annotated as “Macrophage” were selected for further analysis.

scRNAseq integration, dimensionality reduction and clustering: The 3 stable and 3 ALAD samples were downsampled to 1000 Macrophage cells per sample. Then the 3 stable samples were merged and integrated using Seurat SCTransform (54). A similar procedure was applied to the 3 ALAD samples. For each of the integrated datasets (3 stable BAL and 3 ALAD BAL) the following analyses have been performed: Principal component analysis (PCA) was performed using the top 3,000 most variable genes. Then t-distributed stochastic neighbour embedding (tSNE) was performed on the top 30 principal components for visualization of the cells. Subsequently, a graph-based clustering approach using K-nearest neighbor analysis was performed on the top 15 principal components with a resolution of 0.8 for the granularity of the downstream clustering.

Differential gene expression analysis: To measure differential gene expression for each cluster, MAST (56) in Seurat v3 was used to perform differential analysis using normalized data. Similarly DESeq2 (57) was performed for comparing raw RNA expression between individual samples as well as between integrated stable and integrated ALAD samples.

Cell-cell interaction: We employed the CellChat R package (27) to infer intercellular communication based on normalized scRNAseq data. CellChat predicts major signaling inputs and outputs by comparison to cell interactions in the human CellChat Database – which includes cell contact, extracellular matrix-receptor, and secreted signaling data from KEGG and the literature.

Comparison of stable and ALAD datasets: In order to compare the integrated stable and ALAD datasets, we applied the results from differential gene expression analysis and Seurat to the ClusterMap algorithm (58). ClusterMap matched clusters across the two datasets and provided ‘similarity’ as a metric to quantify the quality of the match.

Pathway analysis: Gene ontology (GO) and Gene Set Enrichment Analysis (GSEA) were performed with ClusterProfiler (59) which supports statistical analysis and visualization of functional profiles for genes and gene clusters. In GSEA analysis, 50 hallmark gene sets in MSigDB were used for annotation.

Genotyping scRNAseq samples using a single nucleotide variation (SNV) calling algorithm: To identify donor and recipient cells from scRNA-seq data we used a previously published algorithm, Vireo (60), which was designed to identify cells derived from genetically distinct individuals in synthetically pooled samples. Since transplant samples contain a mixture of cells from genetically distinct individuals, we reasoned that Vireo would enable us to deconvolute donor and recipient cells. Cell Ranger (53) alignments were piled up with the requirement that a called SNV be covered by at least 15 unique cell barcodes, at least 10 unique barcodes supporting the alternative allele, an allele fraction of at least 1%, and to have at least one read with a supporting mismatch outside of the first and last 5 base pairs of the read. We inferred the strand for each potential SNV by requiring that at least 90% of the reads covering the SNV be derived from the same strand. We further filtered these SNVs to remove A>G RNA edits as these are non-genic and not useful for genotype assignments. We identified an SNV as an A>G edit if it was a strand-specific A>G change and present in REDIportal (61) or overlapped an Alu element according to RepeatMasker annotations downloaded from UCSC (62). Genotypes were called by running Vireo v0.3.2 with two genotypes and our SNV dataset with SNVs removed.

After genotyping, we initially observed that there were low-quality genotype calls. Therefore, we developed a second filtering step to remove cells that appeared to have poor genotype assignments. First, we filtered the potential SNVs that were expressed in less than 10 percent of the assigned cells in either genotype. Next, we calculated the summary allele fraction for each genotype/SNV combination summing the (total alternative allele counts) / (total alternative + reference allele counts). Next, for each genotype we selected SNVs that were predictive of the cell's genotype by requiring SNVs to have a receiver operator curve (ROC) AUC >= 0.5 (AUC calculated as |AUC – 0.5| * 2). Next, we selected for SNVs that are homozygous in one genotype and reference in the other as heterozygous SNVs are noisier due to the sparsity of scRNAseq data and ambient RNA contamination. We selected these by requiring the absolute value of the difference between overall allele fractions between the two genotypes be greater than or equal to 0.85. The liberal cut-off below 1.0 is to account for ambient RNA contamination. Next, for each cell we selected the SNVs that are covered by at least one molecule and calculated the average of the absolute value of the difference of each SNV minus the summary SNV for each genotype. The expectation is that each cell should have a low score compared to the assigned genotype and a high score compared to the other genotype. Doublets and unassigned cells are expected to have scores around 0.5 as they are a mix of both genotypes. Finally, we calculated a threshold for each genotype to remove low quality cells with scores higher than the cut-off. The cut-off was set by averaging the difference scores from the cells assigned to the given genotype + 4 median absolute deviations (limited the cut-off to be between 0.15 and 3). We selected all the predicted singlets for each genotype that fell below the cut-off for further analyses.

Identifying a minimum set of SNVs to visualize cell genotype assignments: To find a minimum set of SNVs for each cell we first filtered the list of SNVs to have an AUC of 0.75 (see above) and a mean difference of 0.4 (to include heterogeneous SNVs). Next, we applied a greedy algorithm to find a set of SNVs that covers genotype 1 by selecting each SNV that covers the largest number of uncovered cells from both genotypes. We repeated this for the second genotype to find SNVs that are present in both genotypes.

Functional analysis, flow cytometry, and cytokine array: Fresh BAL samples from 3 ALAD and 4 stable LT recipients were collected and cells were obtained by centrifugation at 400g for 5 minutes at 4°C. BAL cells were cultured at one million cells per millilitre of media (DMEM with 10% FBS) with or without 20ng/ml of LPS. Cultured cells and supernatant were harvested after 16-18 hours and cryopreserved for subsequent experiments. Cells were fixed using 1% formaldehyde before staining with the following surface markers: anti-CD45-BV605 (clone: HI30, BD Biosciences), anti-CD11b-PerCP/Cy5.5 (clone: M1/70), anti-CD163-PE/Dazzle 594 (clone: GHI/61), anti-HLA-DR-Alexa Fluor 647 (clone: L243), and anti-CD32-PE-Cy7 (clone: FUN-2) (all from Biolegend, San Diego, CA), and anti-CD206-APC-Vio 770 (clone: DCN228) (Miltenyi Biotec, Germany). Cells were washed, permeabilized (Cytofix/Cytoperm, BD Biosciences), and intracellular staining was performed with anti-CXCL10-PE (clone: J034D6, Biolegend) and anti-CD68-BV785 (clone: Y1/82A, Biolegend). Cell data were acquired on a BD LSRII flow cytometer and analyzed using Flowjo software. Media supernatants were used to measure released cytokines by BAL cells using a 13-plex bead-based immunoassay and a human macrophage LEGENDplex (Biolegend, San Diego, CA).

**Supplementary figure legends**

**Figure S1. Visualization of quality control metrics for individual samples.** (**A**) Violin plots demonstrating quality control of gene number (nFeature_RNA) between 200 and 5000, unique molecular identifier (UMI) count (nCount_RNA) between 1000 and 30,000 and mitochondrial gene percentage (percent.mt) below 10% of three stable and three ALAD BAL samples. (**B**) Scatter plots showing linear relationship between gene number (Y axis) and UMI count (X axis) for each individual BAL sample.

**Figure S2. The cellular composition of BAL from LT recipients is revealed by single cell RNA sequencing.** (**A**) Data analysis of individual stable and ALAD BAL samples (one sample per row) with reference-based annotation using SingleR algorithm. (**B**) tSNE plots demonstrating between 6 and 10 distinct clusters per sample. (**C**) Distinct gene expression programs depicted in heatmaps of the top differentially expressed genes. (**D**) The total interaction strength (weights) between any two cell groups determined using CellChat is shown using circle plots based on top differentially expressed genes and validated molecular interactions. (**E**) Overall signaling patterns for each cluster is shown in heatmaps where dark green represent higher relative strength of signal.

**Figure S3. Integration cells from 3 stable and 3 ALAD BAL samples using SCTransform**. (**A)** tSNE plot of 3 integrated stable BAL samples. (**B)** tSNE plot of 3 integrated ALAD BAL samples.

**Figure S4. Expression of known macrophage transcripts across AM clusters.** Expression of CD68, CD163, MARCO, MRC1, CD11b, CD16, CD14, CD169, TGM2, CD86, TLR2, FTL, and distinct HLA class II in AMs is shown (darker colour indicates higher relative expression).

**Figure S5. Pathway analysis suggests functional specialization of AMs from stable patients.** **(A)** Pathway analysis performed using GO and GSEA revealed distinct gene programs in the 13 clusters. (**B)** tSNE plot of AMs from stable BAL with cell cycle stage indicated. Distinct clusters of G2M-phase (cluster 10) and S-phase (cluster 11) AMs are seen at lower left. (**C)** Expression of canonical M1- (left) and M2- (right) associated genes in AMs from stable patients. Circle colour reflects average relative gene expression within the cluster, while the size of each circle reflects the percentage of cells within the cluster expressing the indicated gene.

**Figure S6. Predicted pathways through which AM clusters from stable BAL samples communicate with each other.** Significant signaling pathways that are highlighted with red and larger circles have been used to determine potential functionality of AM clusters. (**A**) Significant signaling pathways received by FN1 cluster from other AM clusters (left panel) and sent to other AM clusters (right panel). (**B**) Significant signaling pathways received by PLAC8 cluster from other AM clusters (left panel) and sent to other AM clusters (right panel). (**C**) Significant signaling pathways received by IFI27 cluster from other AM clusters (left panel) and sent to other AM clusters (right panel). (**D**) Significant signaling pathways received by CTNNB1 cluster from other AM clusters (left panel) and sent to other AM clusters (right panel). (**E**) Significant signaling pathways received by SCD cluster from other AM clusters (left panel) and sent to other AM clusters (right panel). (**F**) Significant signaling pathways received by CTSC cluster from other AM clusters (left panel) and sent to other AM clusters (right panel). (**G**) Significant signaling pathways received by MSMO1 cluster from other AM clusters (left panel) and sent to other AM clusters (right panel). (**H**) Significant signaling pathways received by SOD2 cluster from other AM clusters (left panel) and sent to other AM clusters (right panel). (**I**) Significant signaling pathways received by CD32 cluster from other AM clusters (left panel) and sent to other AM clusters (right panel). (**J**) Significant signaling pathways received by EIF cluster from other AM clusters (left panel) and sent to other AM clusters (right panel). (**K**) Significant signaling pathways received by G2/M phase cluster from other AM clusters (left panel) and sent to other AM clusters (right panel). (**L**) Significant signaling pathways received by S phase cluster from other AM clusters (left panel) and sent to other AM clusters (right panel).

**Figure S7. Heatmap showing expression of top differentially expressed genes in 13 AM clusters identified in ALAD BAL samples.** Yellow indicates high relative expression.

**Figure S8. Pathway analysis on ALAD samples using GO and GSEA revealed distinct gene programs in the 13 clusters.**

**Figure S9. A.** tSNE plots showing stable (left) and ALAD (right) samples annotated by outcome of the Clustermap algorithm. In this tSNE format, shared colours and numbers between ALAD and stable samples represent clusters with similar transcriptional profiles. **B.** tSNE plot of AMs from ALAD with cell cycle stage indicated.

**Figure S10. Gating of AMs using flow cytometry.** Live, CD68+HLA-DR+ cells were identified as AMs. In this gate, CD163+CXCL10+ AMs could be identified.

**Figure S11. Heatmap showing expression of top differentially expressed genes in 14 AM clusters identified in CLAD lung tissue samples.** Yellow indicates high relative expression.

**Figure S1**

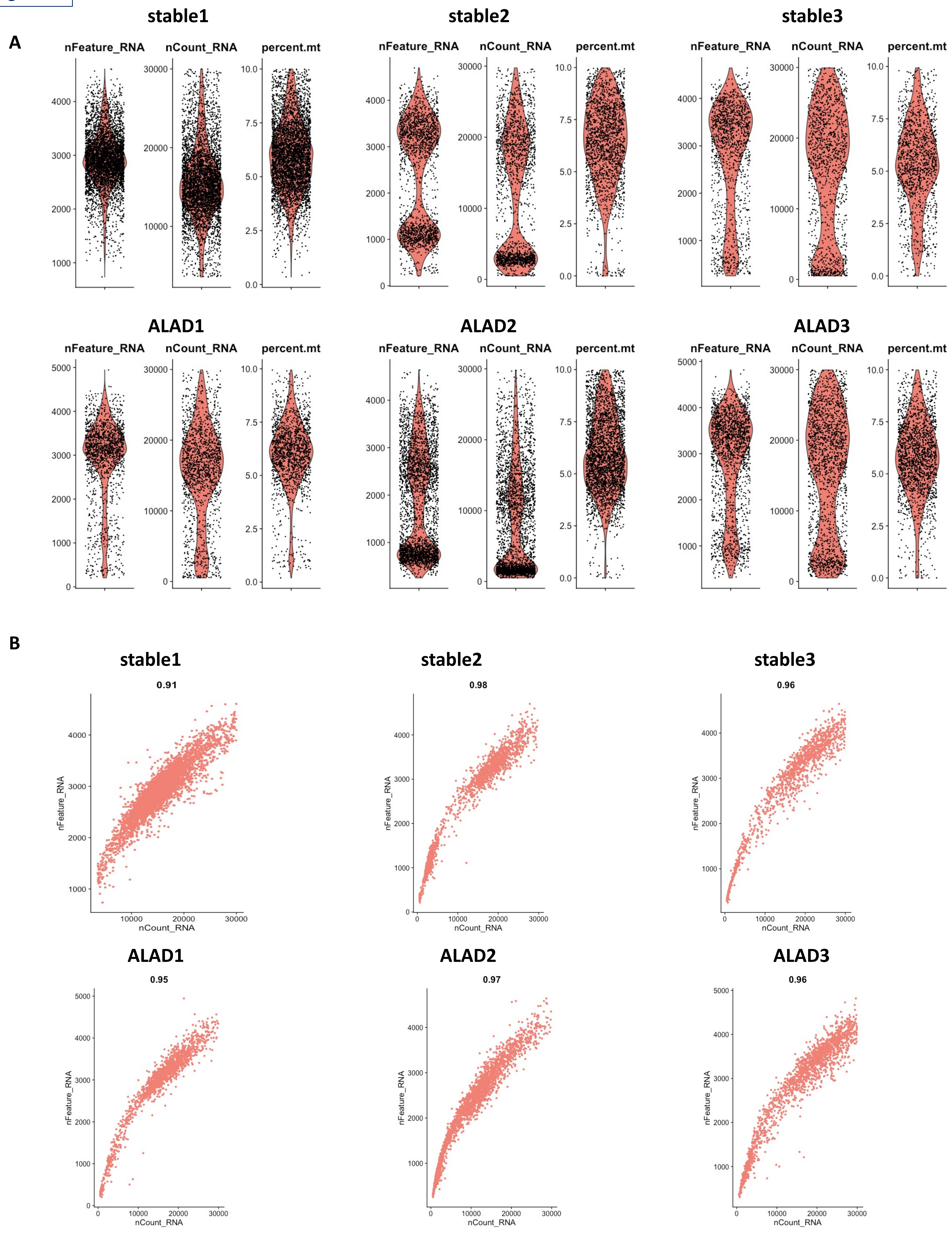
A
stable1
stable2
stable3
ALAD1
ALAD2
ALAD3
nFeature_RNA
nCount_RNA
percent.mt
B
stable1
0.91
stable2
0.98
stable3
0.96
ALAD1
0.95
ALAD2
0.97
ALAD3
0.96
nFeature_RNA
nCount_RNA

Figure S2

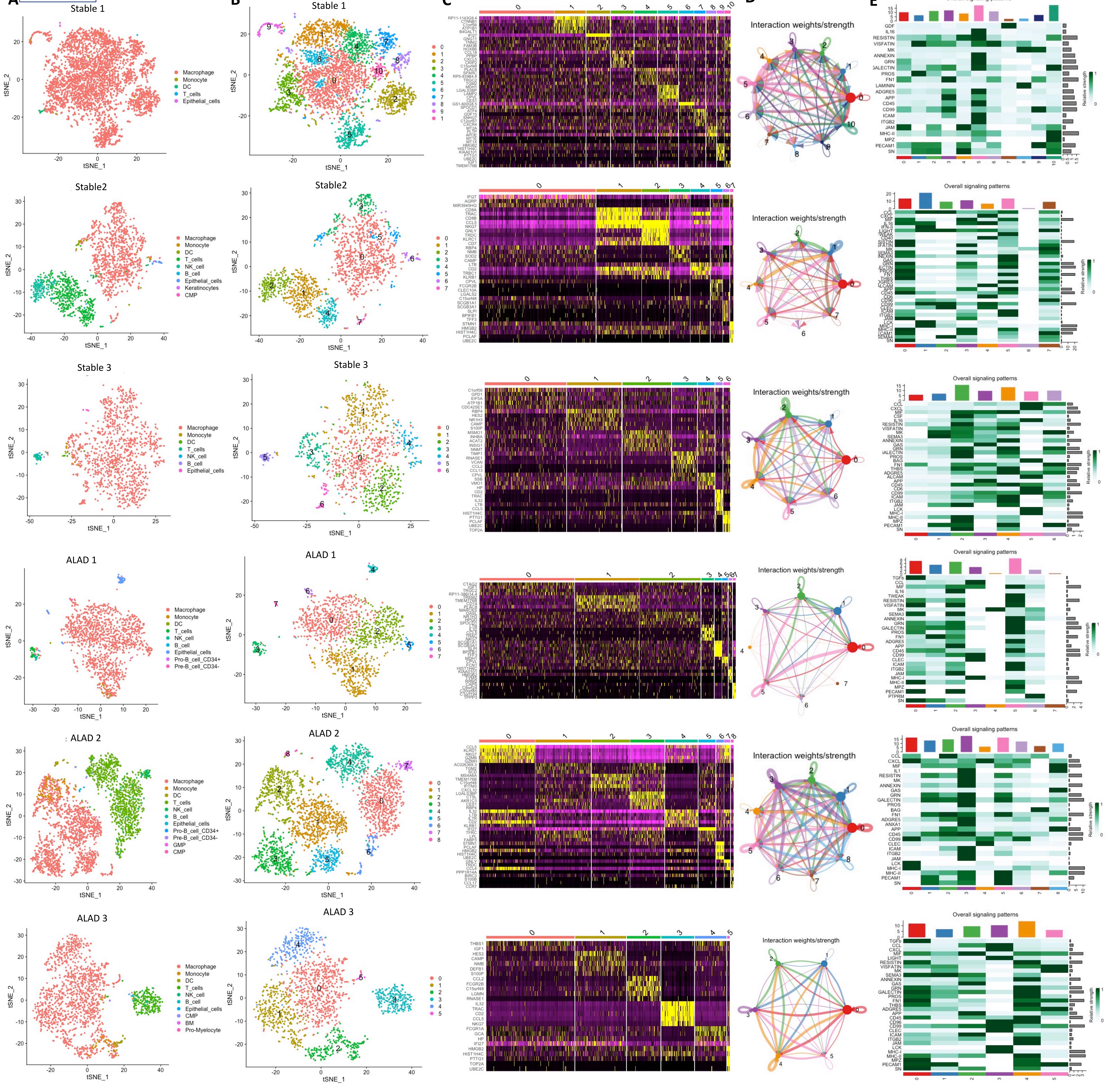
A
B
C
D
E
Stable 1
Stable2
Stable 3
ALAD 1
ALAD 2
ALAD 3
tSNE_1
tSNE_2
Macrophage
Monocyte
DC
T_cells
NK_cell
B_cell
Epithelial_cells
Keratinocytes
CMP
Pro-B_cell_CD34+
Pre-B_cell_CD34-
GMP
BM
Pro-Myelocyte
Interaction weights/strength
Overall signaling patterns
Relative strength

**Figure S3**

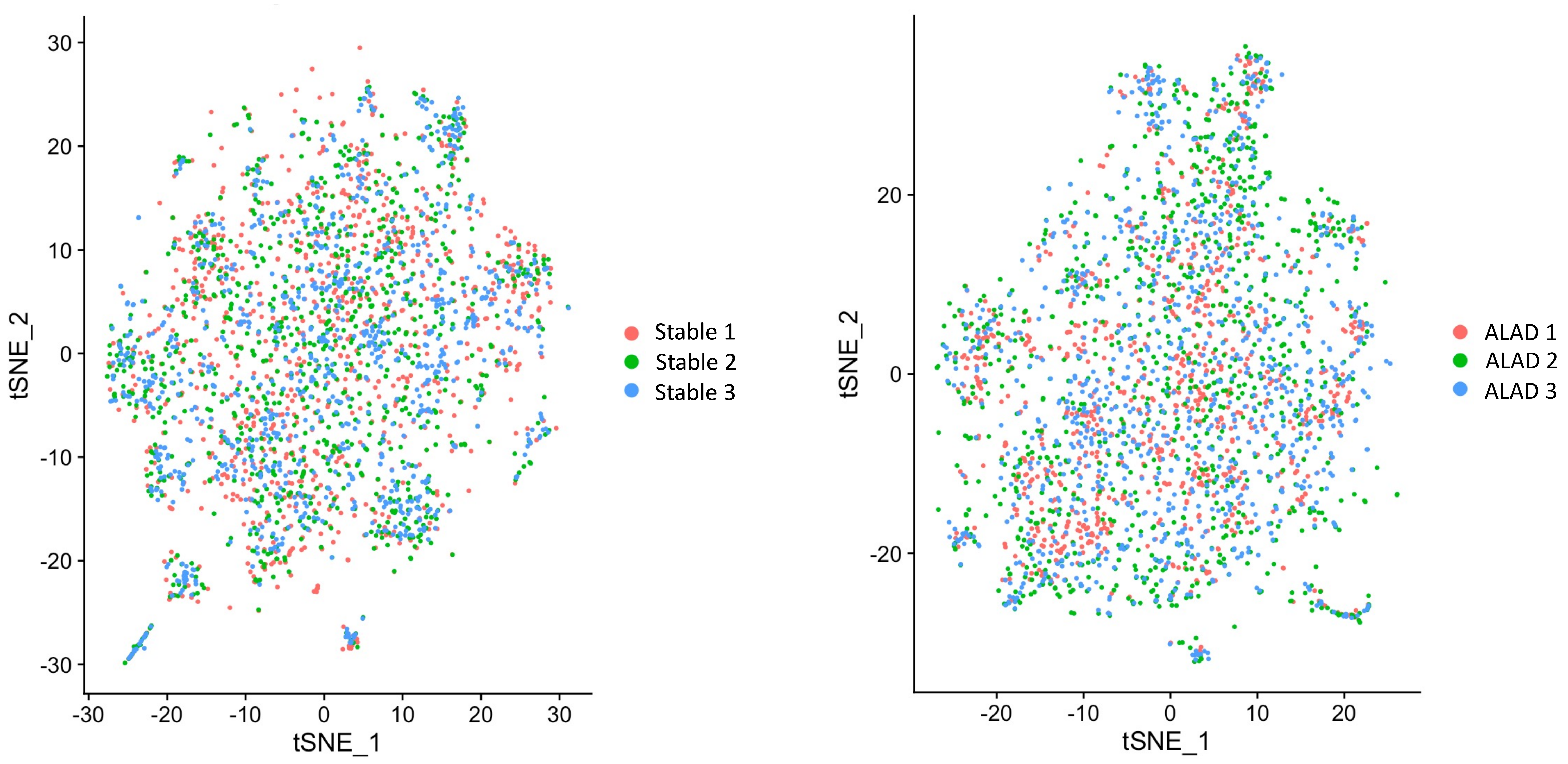
30
20
10
0
-10
-20
-30
tSNE_2
-30
-20
-10
0
10
20
30
tSNE_1
Stable 1
Stable 2
Stable 3
20
0
-20
tSNE_2
-20
-10
0
10
20
tSNE_1
ALAD 1
ALAD 2
ALAD 3

Figure S4

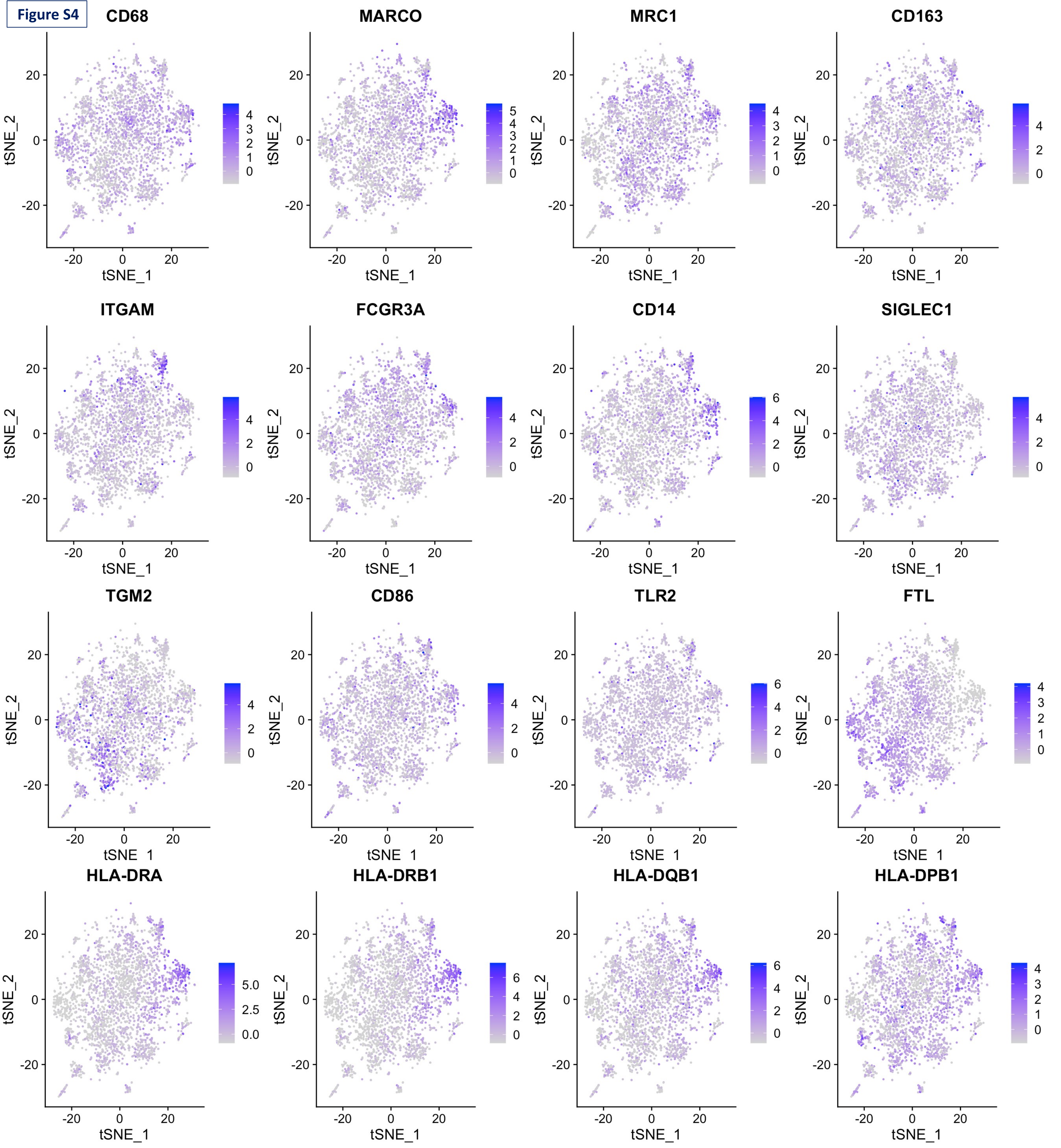
CD68
MARCO
MRC1
CD163
ITGAM
FCGR3A
CD14
SIGLEC1
TGM2
CD86
TLR2
FTL
HLA-DRA
HLA-DRB1
HLA-DQB1
HLA-DPB1
tSNE_1
tSNE_2

**Figure S5**

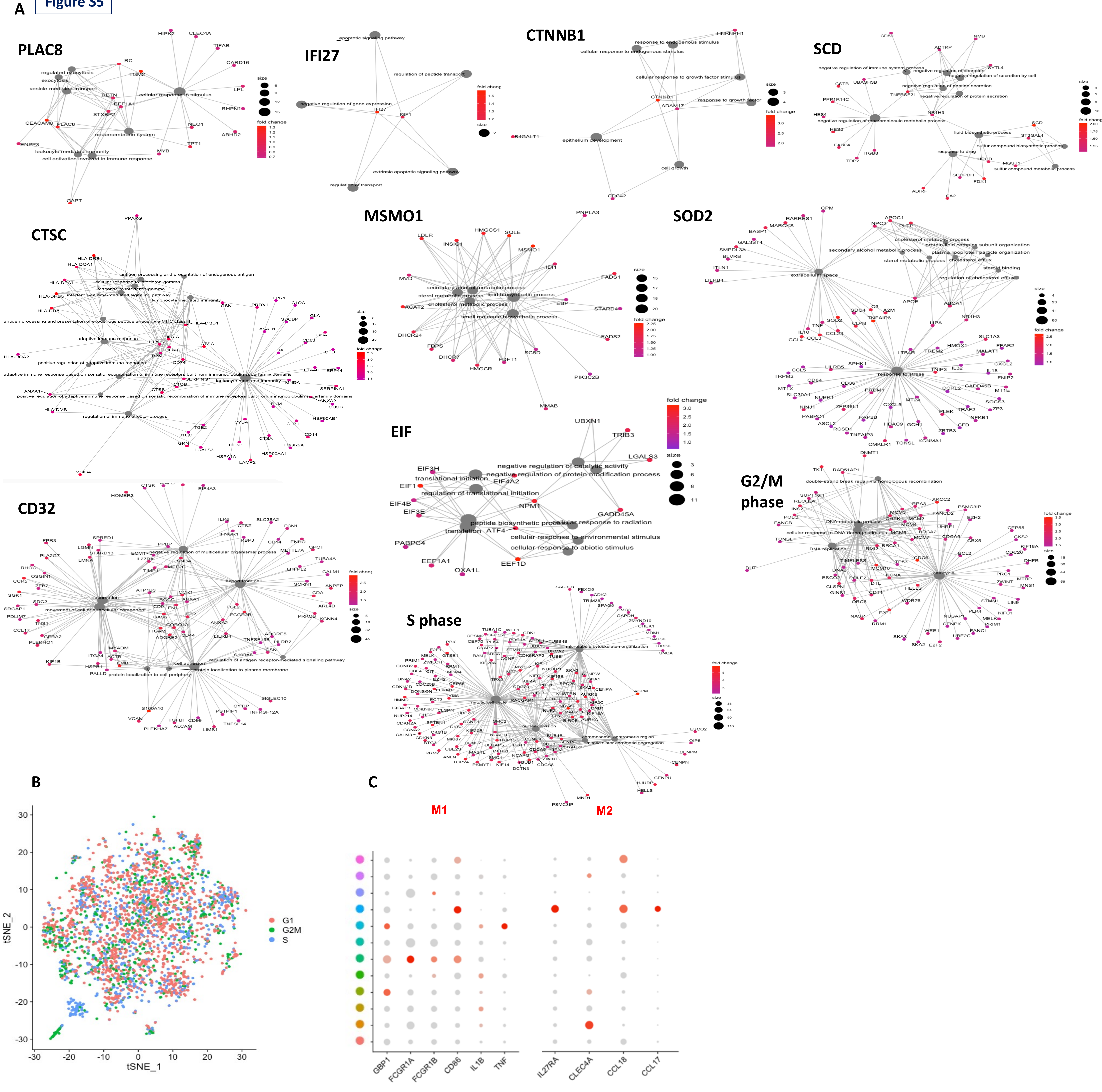

A
PLAC8
IFI27
CTNNB1
SCD
CTSC
MSMO1
SOD2
CD32
EIF
G2/M phase
S phase
size
fold change
B
tSNE_2
tSNE_1
G1
G2M
S
C
M1
M2
GBP1
FCGR1A
FCGR1B
CD86
IL1B
TNF
IL27RA
CLEC4A
CCL18
CCL17

**Figure S6**

**A**

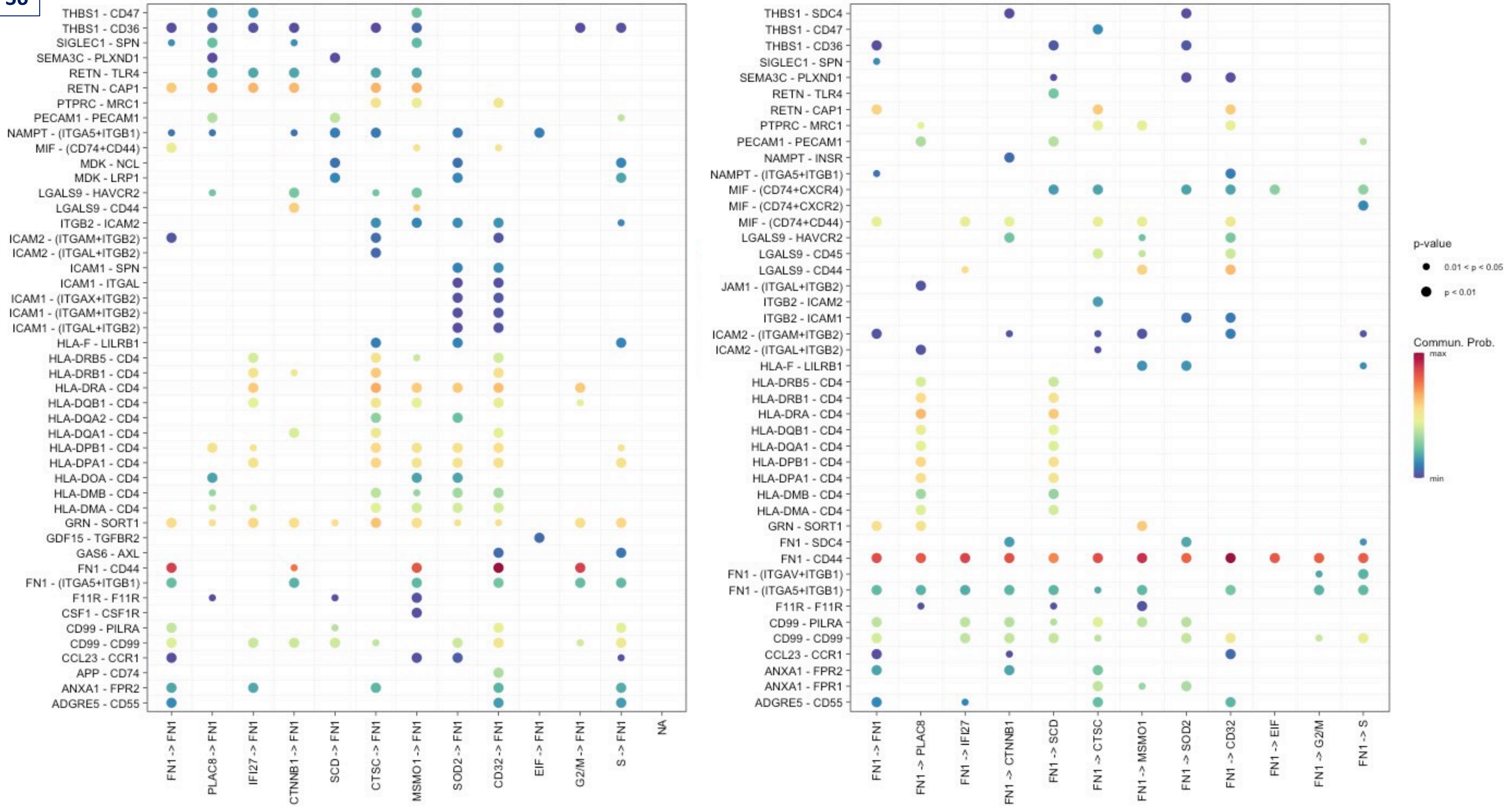
THBS1 - CD47
THBS1 - CD36
SIGLEC1 - SPN
SEMA3C - PLXND1
RETN - TLR4
RETN - CAP1
PTPRC - MRC1
PECAM1 - PECAM1
NAMPT - (ITGA5+ITGB1)
MIF - (CD74+CD44)
MDK - NCL
MDK - LRP1
LGALS9 - HAVCR2
LGALS9 - CD44
ITGB2 - ICAM2
ICAM2 - (ITGAM+ITGB2)
ICAM2 - (ITGAL+ITGB2)
ICAM1 - SPN
ICAM1 - ITGAL
ICAM1 - (ITGAX+ITGB2)
ICAM1 - (ITGAM+ITGB2)
ICAM1 - (ITGAL+ITGB2)
HLA-F - LILRB1
HLA-DRB5 - CD4
HLA-DRB1 - CD4
HLA-DRA - CD4
HLA-DQB1 - CD4
HLA-DQA2 - CD4
HLA-DQA1 - CD4
HLA-DPB1 - CD4
HLA-DPA1 - CD4
HLA-DOA - CD4
HLA-DMB - CD4
HLA-DMA - CD4
GRN - SORT1
GDF15 - TGFBR2
GAS6 - AXL
FN1 - CD44
FN1 - (ITGA5+ITGB1)
F11R - F11R
CSF1 - CSF1R
CD99 - PILRA
CD99 - CD99
CCL23 - CCR1
APP - CD74
ANXA1 - FPR2
ADGRE5 - CD55
FN1 -> FN1
PLAC8 -> FN1
IFI27 -> FN1
CTNNB1 -> FN1
SCD -> FN1
CTSC -> FN1
MSMO1 -> FN1
SOD2 -> FN1
CD32 -> FN1
EIF -> FN1
G2/M -> FN1
S -> FN1
NA
THBS1 - SDC4
THBS1 - CD47
THBS1 - CD36
SIGLEC1 - SPN
SEMA3C - PLXND1
RETN - TLR4
RETN - CAP1
PTPRC - MRC1
PECAM1 - PECAM1
NAMPT - INSR
NAMPT - (ITGA5+ITGB1)
MIF - (CD74+CXCR4)
MIF - (CD74+CXCR2)
MIF - (CD74+CD44)
LGALS9 - HAVCR2
LGALS9 - CD45
LGALS9 - CD44
JAM1 - (ITGAL+ITGB2)
ITGB2 - ICAM2
ITGB2 - ICAM1
ICAM2 - (ITGAM+ITGB2)
ICAM2 - (ITGAL+ITGB2)
HLA-F - LILRB1
HLA-DRB5 - CD4
HLA-DRB1 - CD4
HLA-DRA - CD4
HLA-DQB1 - CD4
HLA-DQA1 - CD4
HLA-DPB1 - CD4
HLA-DPA1 - CD4
HLA-DMB - CD4
HLA-DMA - CD4
GRN - SORT1
FN1 - SDC4
FN1 - CD44
FN1 - (ITGAV+ITGB1)
FN1 - (ITGA5+ITGB1)
F11R - F11R
CD99 - PILRA
CD99 - CD99
CCL23 - CCR1
ANXA1 - FPR2
ANXA1 - FPR1
ADGRE5 - CD55
FN1 -> FN1
FN1 -> PLAC8
FN1 -> IFI27
FN1 -> CTNNB1
FN1 -> SCD
FN1 -> CTSC
FN1 -> MSMO1
FN1 -> SOD2
FN1 -> CD32
FN1 -> EIF
FN1 -> G2/M
FN1 -> S
p-value
0.01 < p < 0.05
p < 0.01
Commun. Prob.
max
min

**B**

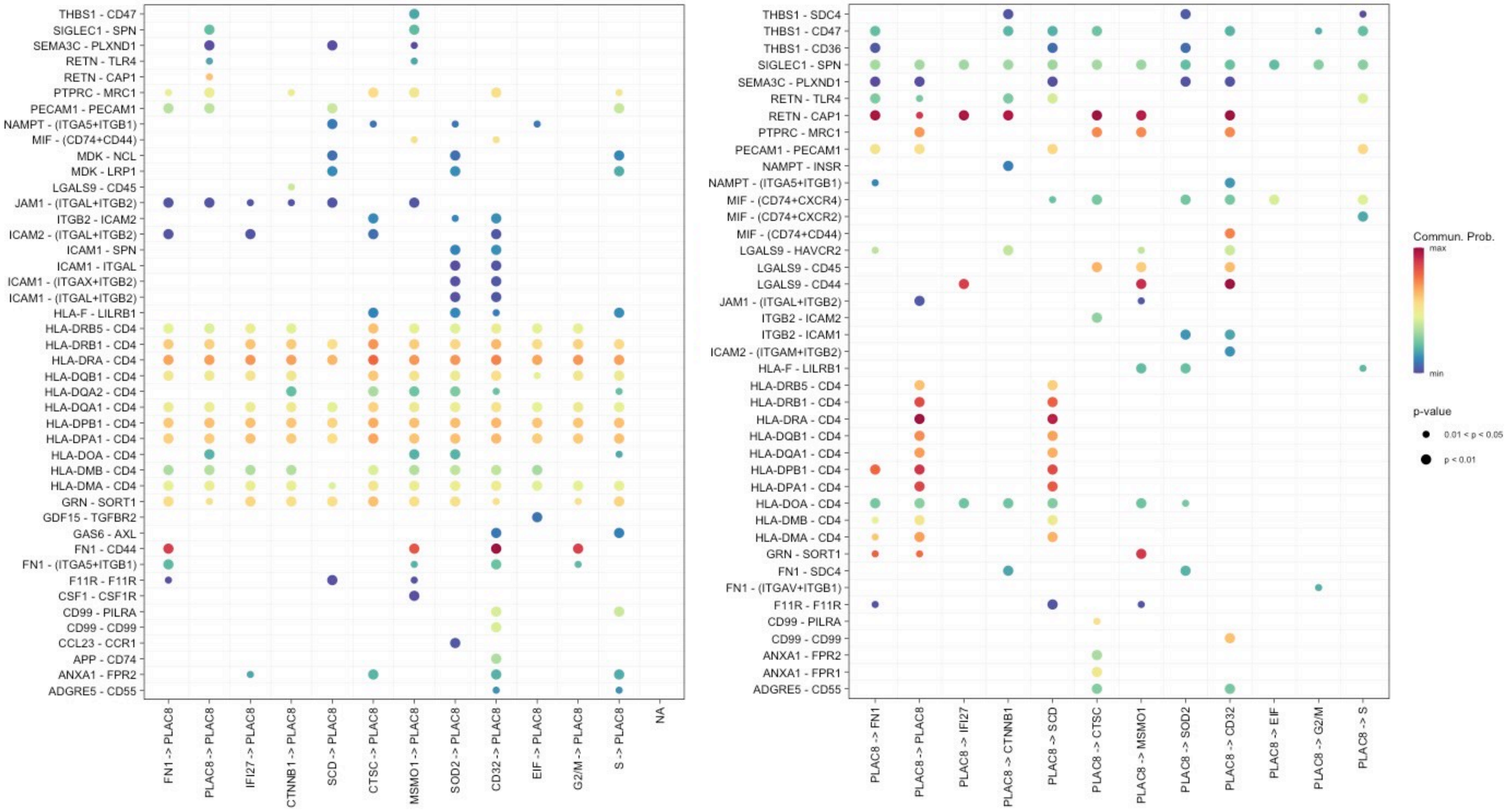
THBS1 - CD47
SIGLEC1 - SPN
SEMA3C - PLXND1
RETN - TLR4
RETN - CAP1
PTPRC - MRC1
PECAM1 - PECAM1
NAMPT - (ITGA5+ITGB1)
MIF - (CD74+CD44)
MDK - NCL
MDK - LRP1
LGALS9 - CD45
JAM1 - (ITGAL+ITGB2)
ITGB2 - ICAM2
ICAM2 - (ITGAL+ITGB2)
ICAM1 - SPN
ICAM1 - ITGAL
ICAM1 - (ITGAX+ITGB2)
ICAM1 - (ITGAL+ITGB2)
HLA-F - LILRB1
HLA-DRB5 - CD4
HLA-DRB1 - CD4
HLA-DRA - CD4
HLA-DQB1 - CD4
HLA-DQA2 - CD4
HLA-DQA1 - CD4
HLA-DPB1 - CD4
HLA-DPA1 - CD4
HLA-DOA - CD4
HLA-DMB - CD4
HLA-DMA - CD4
GRN - SORT1
GDF15 - TGFBR2
GAS6 - AXL
FN1 - CD44
FN1 - (ITGA5+ITGB1)
F11R - F11R
CSF1 - CSF1R
CD99 - PILRA
CD99 - CD99
CCL23 - CCR1
APP - CD74
ANXA1 - FPR2
ADGRE5 - CD55
FN1 -> PLAC8
PLAC8 -> PLAC8
IFI27 -> PLAC8
CTNNB1 -> PLAC8
SCD -> PLAC8
CTSC -> PLAC8
MSMO1 -> PLAC8
SOD2 -> PLAC8
CD32 -> PLAC8
EIF -> PLAC8
G2/M -> PLAC8
S -> PLAC8
NA
THBS1 - SDC4
THBS1 - CD47
THBS1 - CD36
SIGLEC1 - SPN
SEMA3C - PLXND1
RETN - TLR4
RETN - CAP1
PTPRC - MRC1
PECAM1 - PECAM1
NAMPT - INSR
NAMPT - (ITGA5+ITGB1)
MIF - (CD74+CXCR4)
MIF - (CD74+CXCR2)
MIF - (CD74+CD44)
LGALS9 - HAVCR2
LGALS9 - CD45
LGALS9 - CD44
JAM1 - (ITGAL+ITGB2)
ITGB2 - ICAM2
ITGB2 - ICAM1
ICAM2 - (ITGAM+ITGB2)
HLA-F - LILRB1
HLA-DRB5 - CD4
HLA-DRB1 - CD4
HLA-DRA - CD4
HLA-DQB1 - CD4
HLA-DQA1 - CD4
HLA-DPB1 - CD4
HLA-DPA1 - CD4
HLA-DOA - CD4
HLA-DMB - CD4
HLA-DMA - CD4
GRN - SORT1
FN1 - SDC4
FN1 - (ITGAV+ITGB1)
F11R - F11R
CD99 - PILRA
CD99 - CD99
ANXA1 - FPR2
ANXA1 - FPR1
ADGRE5 - CD55
PLAC8 -> FN1
PLAC8 -> PLAC8
PLAC8 -> IFI27
PLAC8 -> CTNNB1
PLAC8 -> SCD
PLAC8 -> CTSC
PLAC8 -> MSMO1
PLAC8 -> SOD2
PLAC8 -> CD32
PLAC8 -> EIF
PLAC8 -> G2/M
PLAC8 -> S
Commun. Prob.
max
min
p-value
0.01 < p < 0.05
p < 0.01

C

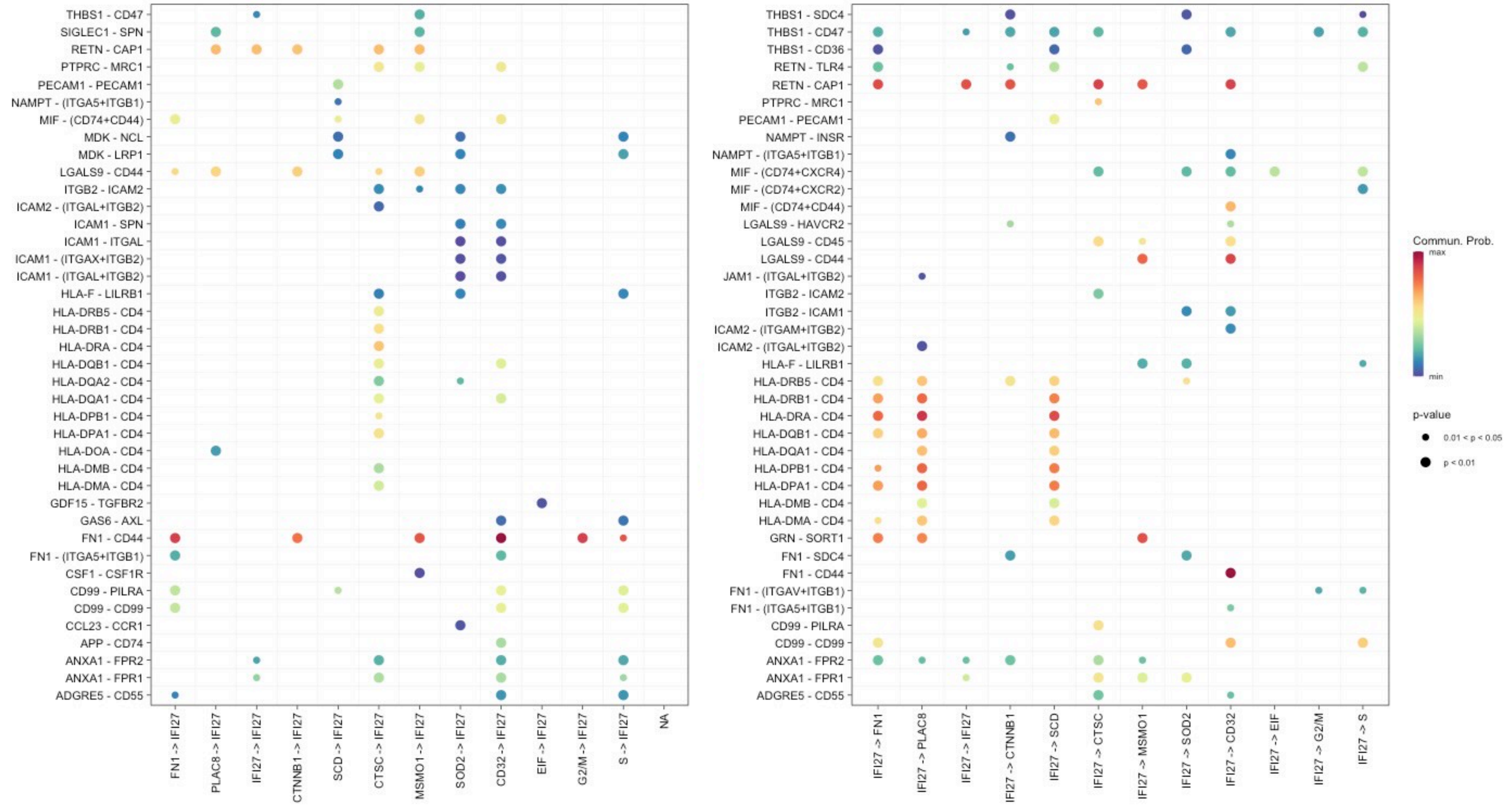
THBS1 - CD47
SIGLEC1 - SPN
RETN - CAP1
PTPRC - MRC1
PECAM1 - PECAM1
NAMPT - (ITGA5+ITGB1)
MIF - (CD74+CD44)
MDK - NCL
MDK - LRP1
LGALS9 - CD44
ITGB2 - ICAM2
ICAM2 - (ITGAL+ITGB2)
ICAM1 - SPN
ICAM1 - ITGAL
ICAM1 - (ITGAX+ITGB2)
ICAM1 - (ITGAL+ITGB2)
HLA-F - LILRB1
HLA-DRB5 - CD4
HLA-DRB1 - CD4
HLA-DRA - CD4
HLA-DQB1 - CD4
HLA-DQA2 - CD4
HLA-DQA1 - CD4
HLA-DPB1 - CD4
HLA-DPA1 - CD4
HLA-DOA - CD4
HLA-DMB - CD4
HLA-DMA - CD4
GDF15 - TGFBR2
GAS6 - AXL
FN1 - CD44
FN1 - (ITGA5+ITGB1)
CSF1 - CSF1R
CD99 - PILRA
CD99 - CD99
CCL23 - CCR1
APP - CD74
ANXA1 - FPR2
ANXA1 - FPR1
ADGRE5 - CD55
FN1 -> IFI27
PLAC8 -> IFI27
IFI27 -> IFI27
CTNNB1 -> IFI27
SCD -> IFI27
CTSC -> IFI27
MSMO1 -> IFI27
SOD2 -> IFI27
CD32 -> IFI27
EIF -> IFI27
G2/M -> IFI27
S -> IFI27
NA
THBS1 - SDC4
THBS1 - CD47
THBS1 - CD36
RETN - TLR4
RETN - CAP1
PTPRC - MRC1
PECAM1 - PECAM1
NAMPT - INSR
NAMPT - (ITGA5+ITGB1)
MIF - (CD74+CXCR4)
MIF - (CD74+CXCR2)
MIF - (CD74+CD44)
LGALS9 - HAVCR2
LGALS9 - CD45
LGALS9 - CD44
JAM1 - (ITGAL+ITGB2)
ITGB2 - ICAM2
ITGB2 - ICAM1
ICAM2 - (ITGAM+ITGB2)
ICAM2 - (ITGAL+ITGB2)
HLA-F - LILRB1
HLA-DRB5 - CD4
HLA-DRB1 - CD4
HLA-DRA - CD4
HLA-DQB1 - CD4
HLA-DQA1 - CD4
HLA-DPB1 - CD4
HLA-DPA1 - CD4
HLA-DMB - CD4
HLA-DMA - CD4
GRN - SORT1
FN1 - SDC4
FN1 - CD44
FN1 - (ITGAV+ITGB1)
FN1 - (ITGA5+ITGB1)
CD99 - PILRA
CD99 - CD99
ANXA1 - FPR2
ANXA1 - FPR1
ADGRE5 - CD55
IFI27 -> FN1
IFI27 -> PLAC8
IFI27 -> IFI27
IFI27 -> CTNNB1
IFI27 -> SCD
IFI27 -> CTSC
IFI27 -> MSMO1
IFI27 -> SOD2
IFI27 -> CD32
IFI27 -> EIF
IFI27 -> G2/M
IFI27 -> S
Commun. Prob.
max
min
p-value
0.01 < p < 0.05
p < 0.01

D

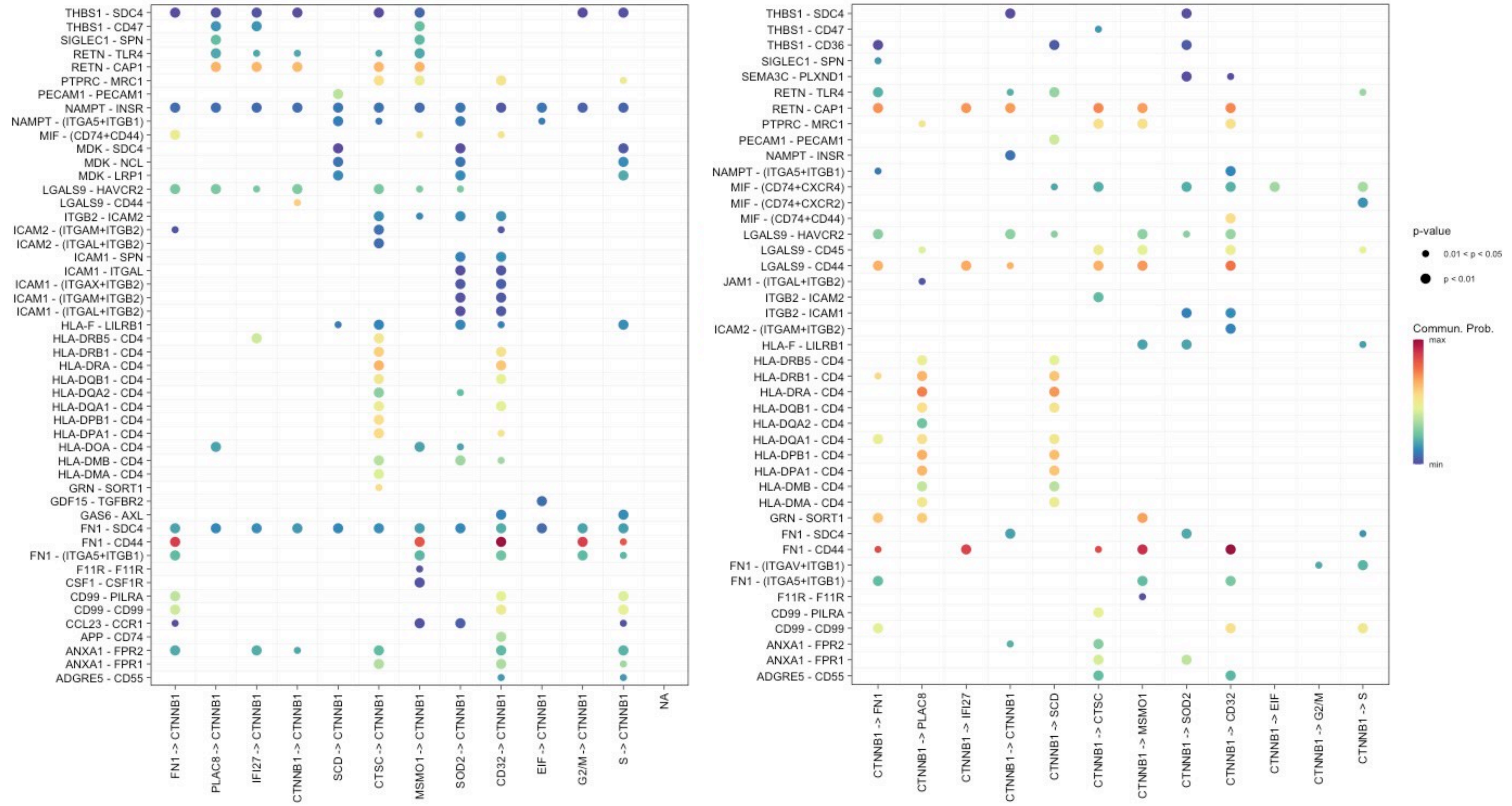
THBS1 - SDC4
THBS1 - CD47
SIGLEC1 - SPN
RETN - TLR4
RETN - CAP1
PTPRC - MRC1
PECAM1 - PECAM1
NAMPT - INSR
NAMPT - (ITGA5+ITGB1)
MIF - (CD74+CD44)
MDK - SDC4
MDK - NCL
MDK - LRP1
LGALS9 - HAVCR2
LGALS9 - CD44
ITGB2 - ICAM2
ICAM2 - (ITGAM+ITGB2)
ICAM2 - (ITGAL+ITGB2)
ICAM1 - SPN
ICAM1 - ITGAL
ICAM1 - (ITGAX+ITGB2)
ICAM1 - (ITGAM+ITGB2)
ICAM1 - (ITGAL+ITGB2)
HLA-F - LILRB1
HLA-DRB5 - CD4
HLA-DRB1 - CD4
HLA-DRA - CD4
HLA-DQB1 - CD4
HLA-DQA2 - CD4
HLA-DQA1 - CD4
HLA-DPB1 - CD4
HLA-DPA1 - CD4
HLA-DOA - CD4
HLA-DMB - CD4
HLA-DMA - CD4
GRN - SORT1
GDF15 - TGFBR2
GAS6 - AXL
FN1 - SDC4
FN1 - CD44
FN1 - (ITGA5+ITGB1)
F11R - F11R
CSF1 - CSF1R
CD99 - PILRA
CD99 - CD99
CCL23 - CCR1
APP - CD74
ANXA1 - FPR2
ANXA1 - FPR1
ADGRE5 - CD55
FN1 -> CTNNB1
PLAC8 -> CTNNB1
IFI27 -> CTNNB1
CTNNB1 -> CTNNB1
SCD -> CTNNB1
CTSC -> CTNNB1
MSMO1 -> CTNNB1
SOD2 -> CTNNB1
CD32 -> CTNNB1
EIF -> CTNNB1
G2/M -> CTNNB1
S -> CTNNB1
NA
THBS1 - SDC4
THBS1 - CD47
THBS1 - CD36
SIGLEC1 - SPN
SEMA3C - PLXND1
RETN - TLR4
RETN - CAP1
PTPRC - MRC1
PECAM1 - PECAM1
NAMPT - INSR
NAMPT - (ITGA5+ITGB1)
MIF - (CD74+CXCR4)
MIF - (CD74+CXCR2)
MIF - (CD74+CD44)
LGALS9 - HAVCR2
LGALS9 - CD45
LGALS9 - CD44
JAM1 - (ITGAL+ITGB2)
ITGB2 - ICAM2
ITGB2 - ICAM1
ICAM2 - (ITGAM+ITGB2)
HLA-F - LILRB1
HLA-DRB5 - CD4
HLA-DRB1 - CD4
HLA-DRA - CD4
HLA-DQB1 - CD4
HLA-DQA2 - CD4
HLA-DQA1 - CD4
HLA-DPB1 - CD4
HLA-DPA1 - CD4
HLA-DMB - CD4
HLA-DMA - CD4
GRN - SORT1
FN1 - SDC4
FN1 - CD44
FN1 - (ITGAV+ITGB1)
FN1 - (ITGA5+ITGB1)
F11R - F11R
CD99 - PILRA
CD99 - CD99
ANXA1 - FPR2
ANXA1 - FPR1
ADGRE5 - CD55
CTNNB1 -> FN1
CTNNB1 -> PLAC8
CTNNB1 -> IFI27
CTNNB1 -> CTNNB1
CTNNB1 -> SCD
CTNNB1 -> CTSC
CTNNB1 -> MSMO1
CTNNB1 -> SOD2
CTNNB1 -> CD32
CTNNB1 -> EIF
CTNNB1 -> G2/M
CTNNB1 -> S
p-value
0.01 < p < 0.05
p < 0.01
Commun. Prob.
max
min

E

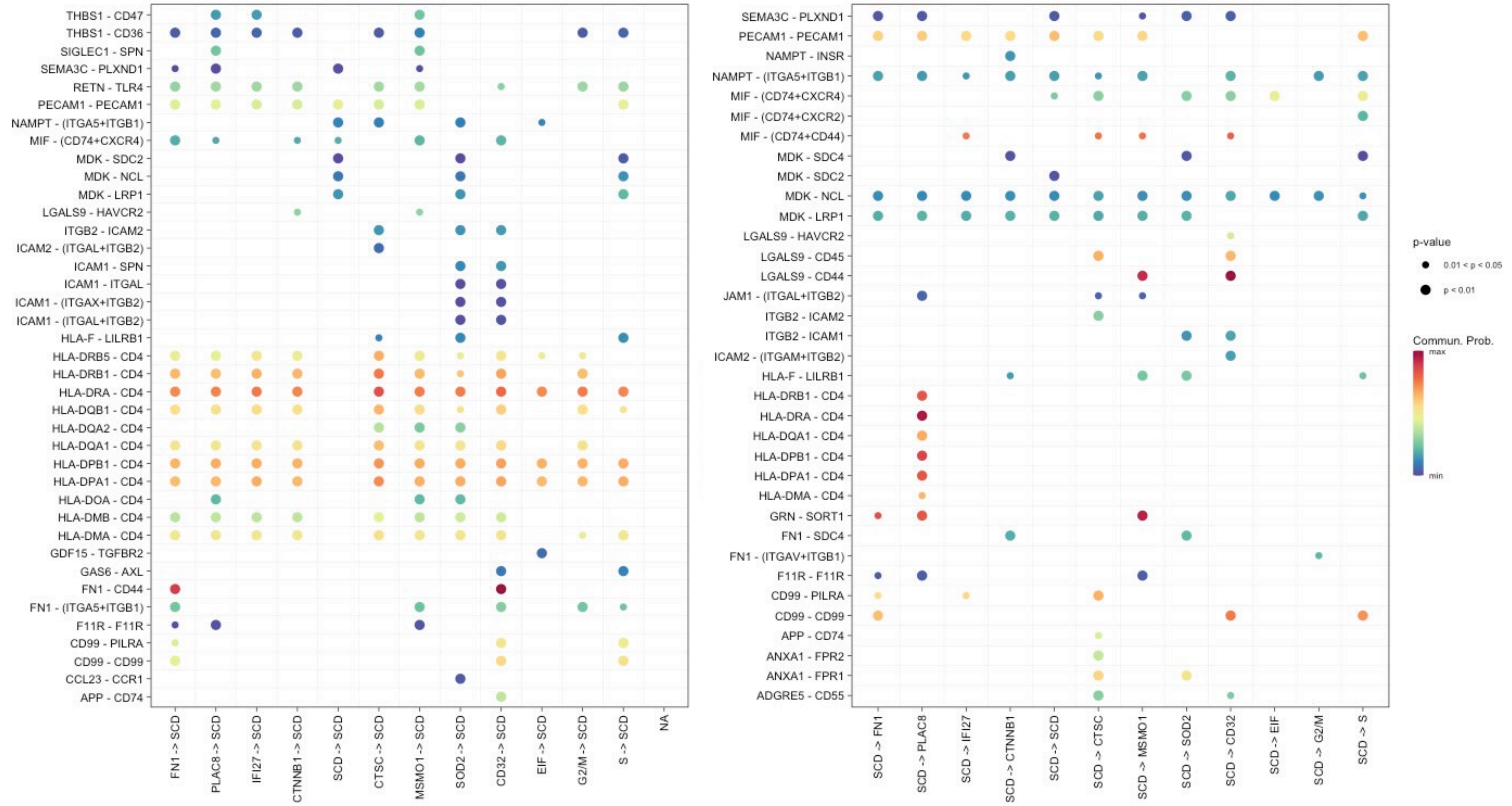
THBS1 - CD47
THBS1 - CD36
SIGLEC1 - SPN
SEMA3C - PLXND1
RETN - TLR4
PECAM1 - PECAM1
NAMPT - (ITGA5+ITGB1)
MIF - (CD74+CXCR4)
MDK - SDC2
MDK - NCL
MDK - LRP1
LGALS9 - HAVCR2
ITGB2 - ICAM2
ICAM2 - (ITGAL+ITGB2)
ICAM1 - SPN
ICAM1 - ITGAL
ICAM1 - (ITGAX+ITGB2)
ICAM1 - (ITGAL+ITGB2)
HLA-F - LILRB1
HLA-DRB5 - CD4
HLA-DRB1 - CD4
HLA-DRA - CD4
HLA-DQB1 - CD4
HLA-DQA2 - CD4
HLA-DQA1 - CD4
HLA-DPB1 - CD4
HLA-DPA1 - CD4
HLA-DOA - CD4
HLA-DMB - CD4
HLA-DMA - CD4
GDF15 - TGFBR2
GAS6 - AXL
FN1 - CD44
FN1 - (ITGA5+ITGB1)
F11R - F11R
CD99 - PILRA
CD99 - CD99
CCL23 - CCR1
APP - CD74
FN1 -> SCD
PLAC8 -> SCD
IFI27 -> SCD
CTNNB1 -> SCD
SCD -> SCD
CTSC -> SCD
MSMO1 -> SCD
SOD2 -> SCD
CD32 -> SCD
EIF -> SCD
G2/M -> SCD
S -> SCD
NA
SEMA3C - PLXND1
PECAM1 - PECAM1
NAMPT - INSR
NAMPT - (ITGA5+ITGB1)
MIF - (CD74+CXCR4)
MIF - (CD74+CXCR2)
MIF - (CD74+CD44)
MDK - SDC4
MDK - SDC2
MDK - NCL
MDK - LRP1
LGALS9 - HAVCR2
LGALS9 - CD45
LGALS9 - CD44
JAM1 - (ITGAL+ITGB2)
ITGB2 - ICAM2
ITGB2 - ICAM1
ICAM2 - (ITGAM+ITGB2)
HLA-F - LILRB1
HLA-DRB1 - CD4
HLA-DRA - CD4
HLA-DQA1 - CD4
HLA-DPB1 - CD4
HLA-DPA1 - CD4
HLA-DMA - CD4
GRN - SORT1
FN1 - SDC4
FN1 - (ITGAV+ITGB1)
F11R - F11R
CD99 - PILRA
CD99 - CD99
APP - CD74
ANXA1 - FPR2
ANXA1 - FPR1
ADGRE5 - CD55
SCD -> FN1
SCD -> PLAC8
SCD -> IFI27
SCD -> CTNNB1
SCD -> SCD
SCD -> CTSC
SCD -> MSMO1
SCD -> SOD2
SCD -> CD32
SCD -> EIF
SCD -> G2/M
SCD -> S
p-value
0.01 < p < 0.05
p < 0.01
Commun. Prob.
max
min

F

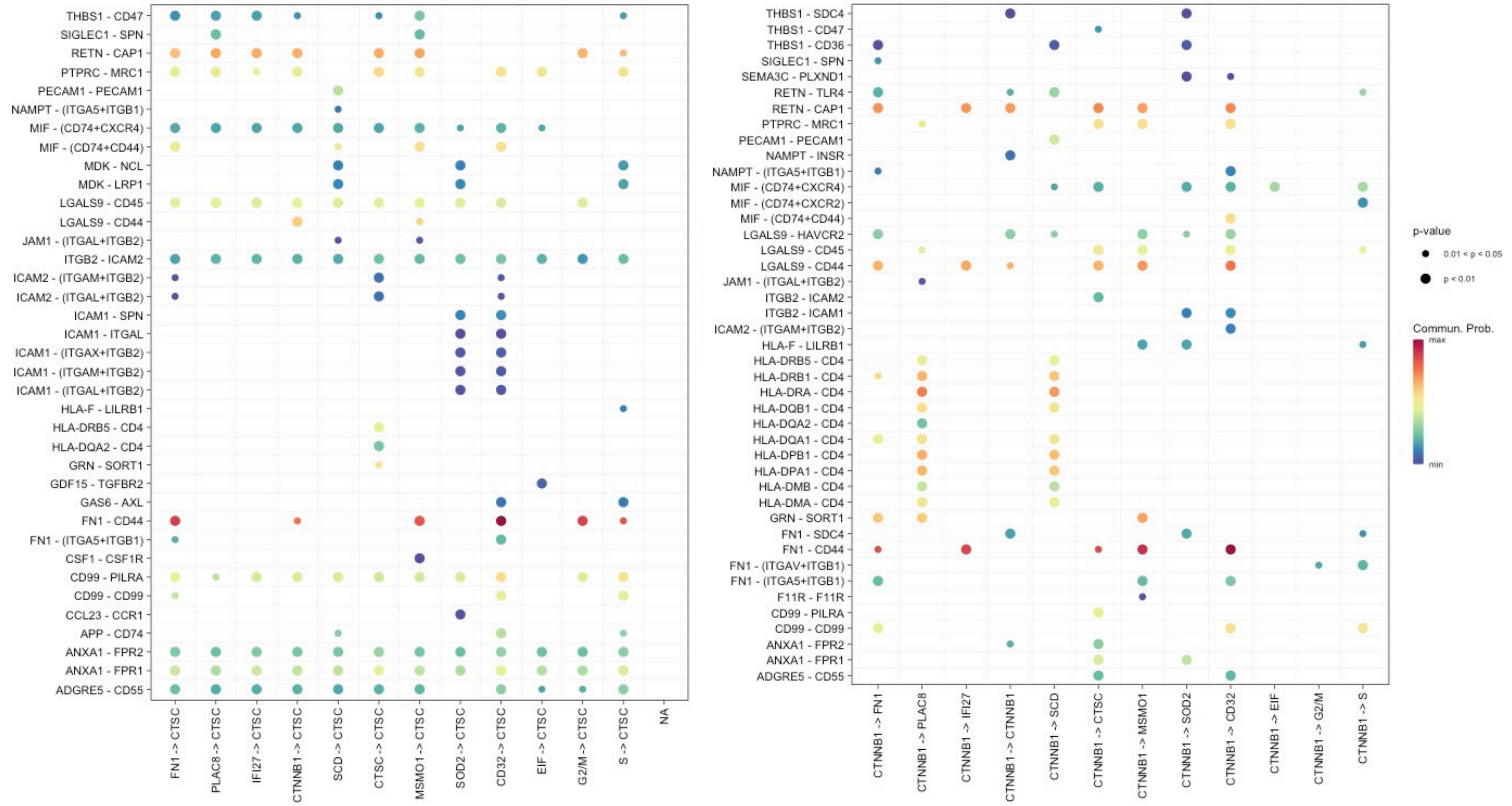
THBS1 - CD47
SIGLEC1 - SPN
RETN - CAP1
PTPRC - MRC1
PECAM1 - PECAM1
NAMPT - (ITGA5+ITGB1)
MIF - (CD74+CXCR4)
MIF - (CD74+CD44)
MDK - NCL
MDK - LRP1
LGALS9 - CD45
LGALS9 - CD44
JAM1 - (ITGAL+ITGB2)
ITGB2 - ICAM2
ICAM2 - (ITGAM+ITGB2)
ICAM2 - (ITGAL+ITGB2)
ICAM1 - SPN
ICAM1 - ITGAL
ICAM1 - (ITGAX+ITGB2)
ICAM1 - (ITGAM+ITGB2)
ICAM1 - (ITGAL+ITGB2)
HLA-F - LILRB1
HLA-DRB5 - CD4
HLA-DQA2 - CD4
GRN - SORT1
GDF15 - TGFBR2
GAS6 - AXL
FN1 - CD44
FN1 - (ITGA5+ITGB1)
CSF1 - CSF1R
CD99 - PILRA
CD99 - CD99
CCL23 - CCR1
APP - CD74
ANXA1 - FPR2
ANXA1 - FPR1
ADGRE5 - CD55
FN1 -> CTSC
PLAC8 -> CTSC
IFI27 -> CTSC
CTNNB1 -> CTSC
SCD -> CTSC
CTSC -> CTSC
MSMO1 -> CTSC
SOD2 -> CTSC
CD32 -> CTSC
EIF -> CTSC
G2/M -> CTSC
S -> CTSC
NA
THBS1 - SDC4
THBS1 - CD47
THBS1 - CD36
SIGLEC1 - SPN
SEMA3C - PLXND1
RETN - TLR4
RETN - CAP1
PTPRC - MRC1
PECAM1 - PECAM1
NAMPT - INSR
NAMPT - (ITGA5+ITGB1)
MIF - (CD74+CXCR4)
MIF - (CD74+CXCR2)
MIF - (CD74+CD44)
LGALS9 - HAVCR2
LGALS9 - CD45
LGALS9 - CD44
JAM1 - (ITGAL+ITGB2)
ITGB2 - ICAM2
ITGB2 - ICAM1
ICAM2 - (ITGAM+ITGB2)
HLA-F - LILRB1
HLA-DRB5 - CD4
HLA-DRB1 - CD4
HLA-DRA - CD4
HLA-DQB1 - CD4
HLA-DQA2 - CD4
HLA-DQA1 - CD4
HLA-DPB1 - CD4
HLA-DPA1 - CD4
HLA-DMB - CD4
HLA-DMA - CD4
GRN - SORT1
FN1 - SDC4
FN1 - CD44
FN1 - (ITGAV+ITGB1)
FN1 - (ITGA5+ITGB1)
F11R - F11R
CD99 - PILRA
CD99 - CD99
ANXA1 - FPR2
ANXA1 - FPR1
ADGRE5 - CD55
CTNNB1 -> FN1
CTNNB1 -> PLAC8
CTNNB1 -> IFI27
CTNNB1 -> CTNNB1
CTNNB1 -> SCD
CTNNB1 -> CTSC
CTNNB1 -> MSMO1
CTNNB1 -> SOD2
CTNNB1 -> CD32
CTNNB1 -> EIF
CTNNB1 -> G2/M
CTNNB1 -> S
p-value
0.01 < p < 0.05
p < 0.01
Commun. Prob.
max
min

G

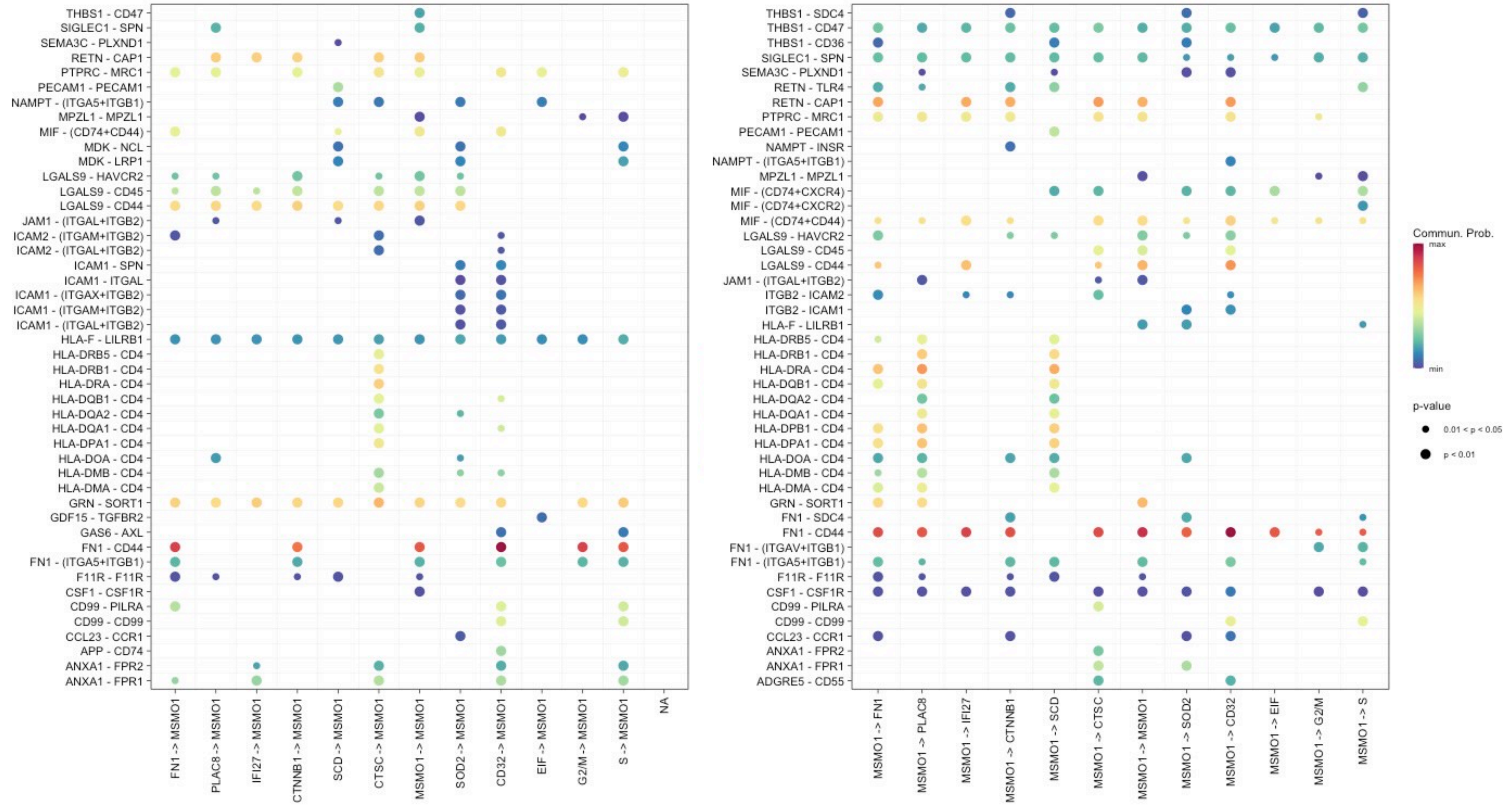
THBS1 - CD47
SIGLEC1 - SPN
SEMA3C - PLXND1
RETN - CAP1
PTPRC - MRC1
PECAM1 - PECAM1
NAMPT - (ITGA5+ITGB1)
MPZL1 - MPZL1
MIF - (CD74+CD44)
MDK - NCL
MDK - LRP1
LGALS9 - HAVCR2
LGALS9 - CD45
LGALS9 - CD44
JAM1 - (ITGAL+ITGB2)
ICAM2 - (ITGAM+ITGB2)
ICAM2 - (ITGAL+ITGB2)
ICAM1 - SPN
ICAM1 - ITGAL
ICAM1 - (ITGAX+ITGB2)
ICAM1 - (ITGAM+ITGB2)
ICAM1 - (ITGAL+ITGB2)
HLA-F - LILRB1
HLA-DRB5 - CD4
HLA-DRB1 - CD4
HLA-DRA - CD4
HLA-DQB1 - CD4
HLA-DQA2 - CD4
HLA-DQA1 - CD4
HLA-DPA1 - CD4
HLA-DOA - CD4
HLA-DMB - CD4
HLA-DMA - CD4
GRN - SORT1
GDF15 - TGFBR2
GAS6 - AXL
FN1 - CD44
FN1 - (ITGA5+ITGB1)
F11R - F11R
CSF1 - CSF1R
CD99 - PILRA
CD99 - CD99
CCL23 - CCR1
APP - CD74
ANXA1 - FPR2
ANXA1 - FPR1
FN1 -> MSMO1
PLAC8 -> MSMO1
IFI27 -> MSMO1
CTNNB1 -> MSMO1
SCD -> MSMO1
CTSC -> MSMO1
MSMO1 -> MSMO1
SOD2 -> MSMO1
CD32 -> MSMO1
EIF -> MSMO1
G2/M -> MSMO1
S -> MSMO1
NA
THBS1 - SDC4
THBS1 - CD47
THBS1 - CD36
SIGLEC1 - SPN
SEMA3C - PLXND1
RETN - TLR4
RETN - CAP1
PTPRC - MRC1
PECAM1 - PECAM1
NAMPT - INSR
NAMPT - (ITGA5+ITGB1)
MPZL1 - MPZL1
MIF - (CD74+CXCR4)
MIF - (CD74+CXCR2)
MIF - (CD74+CD44)
LGALS9 - HAVCR2
LGALS9 - CD45
LGALS9 - CD44
JAM1 - (ITGAL+ITGB2)
ITGB2 - ICAM2
ITGB2 - ICAM1
HLA-F - LILRB1
HLA-DRB5 - CD4
HLA-DRB1 - CD4
HLA-DRA - CD4
HLA-DQB1 - CD4
HLA-DQA2 - CD4
HLA-DQA1 - CD4
HLA-DPB1 - CD4
HLA-DPA1 - CD4
HLA-DOA - CD4
HLA-DMB - CD4
HLA-DMA - CD4
GRN - SORT1
FN1 - SDC4
FN1 - CD44
FN1 - (ITGAV+ITGB1)
FN1 - (ITGA5+ITGB1)
F11R - F11R
CSF1 - CSF1R
CD99 - PILRA
CD99 - CD99
CCL23 - CCR1
ANXA1 - FPR2
ANXA1 - FPR1
ADGRE5 - CD55
MSMO1 -> FN1
MSMO1 -> PLAC8
MSMO1 -> IFI27
MSMO1 -> CTNNB1
MSMO1 -> SCD
MSMO1 -> CTSC
MSMO1 -> MSMO1
MSMO1 -> SOD2
MSMO1 -> CD32
MSMO1 -> EIF
MSMO1 -> G2/M
MSMO1 -> S
Commun. Prob.
max
min
p-value
0.01 < p < 0.05
p < 0.01

H

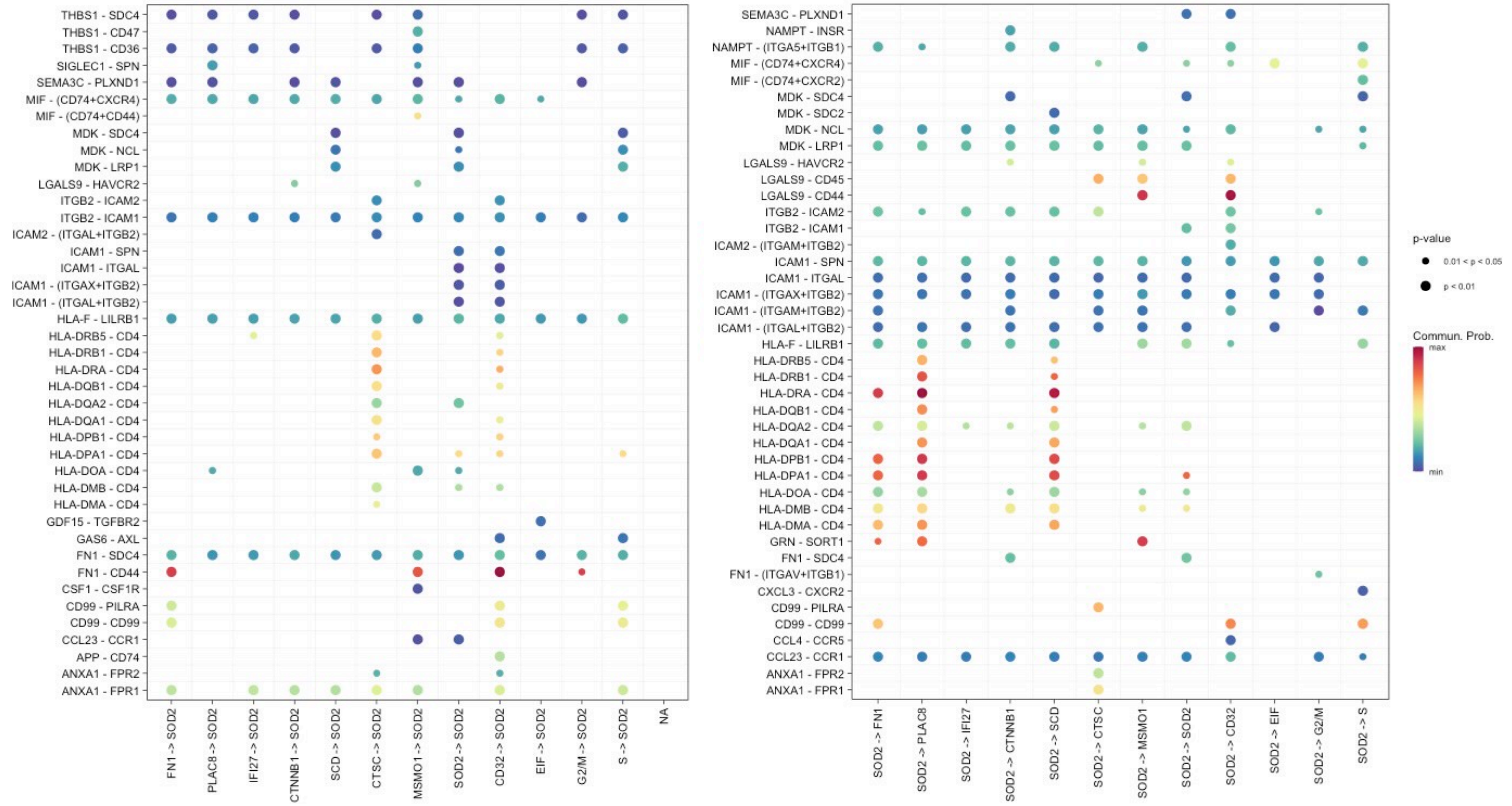
THBS1 - SDC4
THBS1 - CD47
THBS1 - CD36
SIGLEC1 - SPN
SEMA3C - PLXND1
MIF - (CD74+CXCR4)
MIF - (CD74+CD44)
MDK - SDC4
MDK - NCL
MDK - LRP1
LGALS9 - HAVCR2
ITGB2 - ICAM2
ITGB2 - ICAM1
ICAM2 - (ITGAL+ITGB2)
ICAM1 - SPN
ICAM1 - ITGAL
ICAM1 - (ITGAX+ITGB2)
ICAM1 - (ITGAL+ITGB2)
HLA-F - LILRB1
HLA-DRB5 - CD4
HLA-DRB1 - CD4
HLA-DRA - CD4
HLA-DQB1 - CD4
HLA-DQA2 - CD4
HLA-DQA1 - CD4
HLA-DPB1 - CD4
HLA-DPA1 - CD4
HLA-DOA - CD4
HLA-DMB - CD4
HLA-DMA - CD4
GDF15 - TGFBR2
GAS6 - AXL
FN1 - SDC4
FN1 - CD44
CSF1 - CSF1R
CD99 - PILRA
CD99 - CD99
CCL23 - CCR1
APP - CD74
ANXA1 - FPR2
ANXA1 - FPR1
FN1 -> SOD2
PLAC8 -> SOD2
IFI27 -> SOD2
CTNNB1 -> SOD2
SCD -> SOD2
CTSC -> SOD2
MSMO1 -> SOD2
SOD2 -> SOD2
CD32 -> SOD2
EIF -> SOD2
G2/M -> SOD2
S -> SOD2
NA
SEMA3C - PLXND1
NAMPT - INSR
NAMPT - (ITGA5+ITGB1)
MIF - (CD74+CXCR4)
MIF - (CD74+CXCR2)
MDK - SDC4
MDK - SDC2
MDK - NCL
MDK - LRP1
LGALS9 - HAVCR2
LGALS9 - CD45
LGALS9 - CD44
ITGB2 - ICAM2
ITGB2 - ICAM1
ICAM2 - (ITGAM+ITGB2)
ICAM1 - SPN
ICAM1 - ITGAL
ICAM1 - (ITGAX+ITGB2)
ICAM1 - (ITGAM+ITGB2)
ICAM1 - (ITGAL+ITGB2)
HLA-F - LILRB1
HLA-DRB5 - CD4
HLA-DRB1 - CD4
HLA-DRA - CD4
HLA-DQB1 - CD4
HLA-DQA2 - CD4
HLA-DQA1 - CD4
HLA-DPB1 - CD4
HLA-DPA1 - CD4
HLA-DOA - CD4
HLA-DMB - CD4
HLA-DMA - CD4
GRN - SORT1
FN1 - SDC4
FN1 - (ITGAV+ITGB1)
CXCL3 - CXCR2
CD99 - PILRA
CD99 - CD99
CCL4 - CCR5
CCL23 - CCR1
ANXA1 - FPR2
ANXA1 - FPR1
SOD2 -> FN1
SOD2 -> PLAC8
SOD2 -> IFI27
SOD2 -> CTNNB1
SOD2 -> SCD
SOD2 -> CTSC
SOD2 -> MSMO1
SOD2 -> SOD2
SOD2 -> CD32
SOD2 -> EIF
SOD2 -> G2/M
SOD2 -> S
p-value
0.01 < p < 0.05
p < 0.01
Commun. Prob.
max
min

I

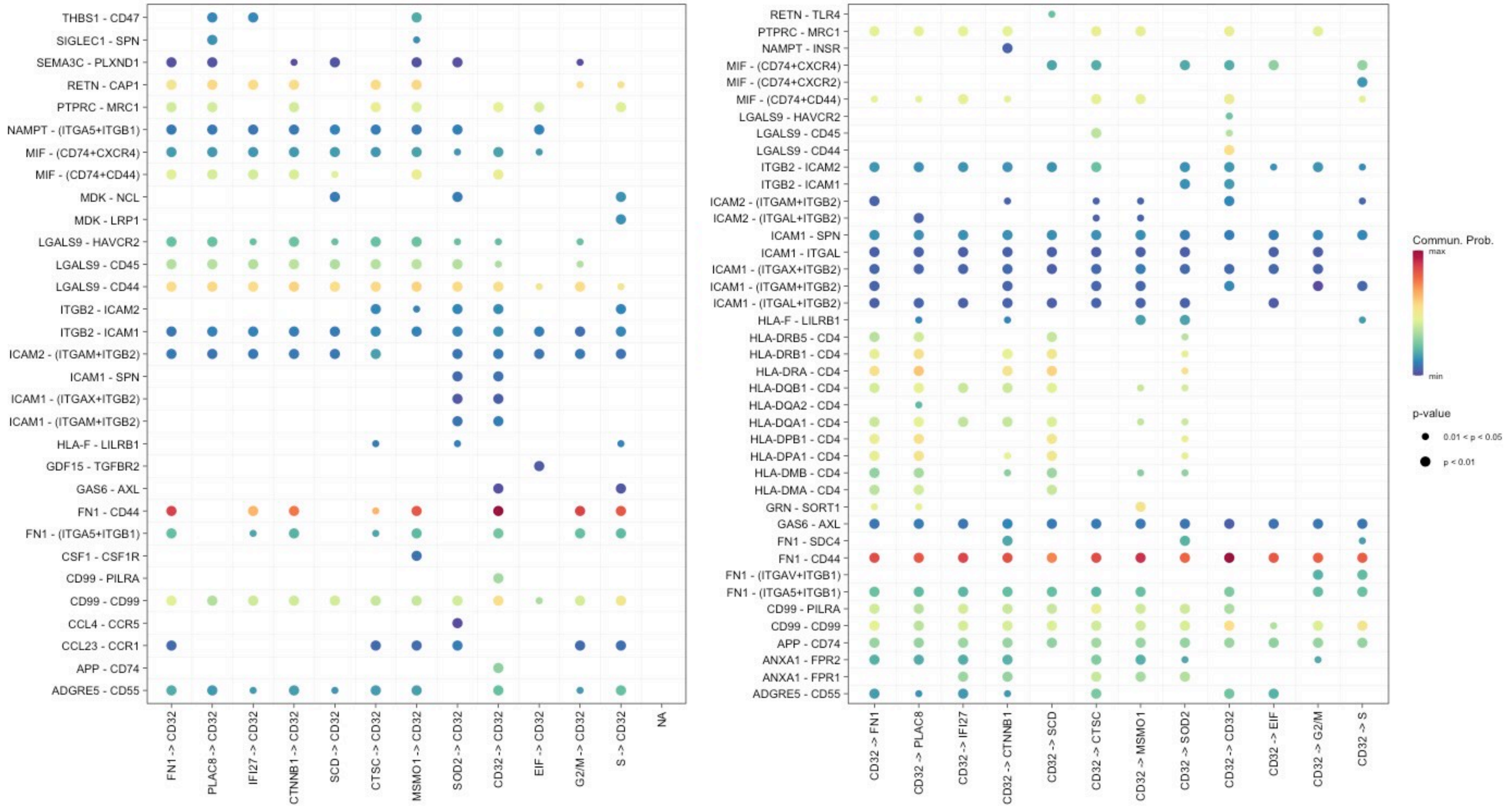
THBS1 - CD47
SIGLEC1 - SPN
SEMA3C - PLXND1
RETN - CAP1
PTPRC - MRC1
NAMPT - (ITGA5+ITGB1)
MIF - (CD74+CXCR4)
MIF - (CD74+CD44)
MDK - NCL
MDK - LRP1
LGALS9 - HAVCR2
LGALS9 - CD45
LGALS9 - CD44
ITGB2 - ICAM2
ITGB2 - ICAM1
ICAM2 - (ITGAM+ITGB2)
ICAM1 - SPN
ICAM1 - (ITGAX+ITGB2)
ICAM1 - (ITGAM+ITGB2)
HLA-F - LILRB1
GDF15 - TGFBR2
GAS6 - AXL
FN1 - CD44
FN1 - (ITGA5+ITGB1)
CSF1 - CSF1R
CD99 - PILRA
CD99 - CD99
CCL4 - CCR5
CCL23 - CCR1
APP - CD74
ADGRE5 - CD55
FN1 -> CD32
PLAC8 -> CD32
IFI27 -> CD32
CTNNB1 -> CD32
SCD -> CD32
CTSC -> CD32
MSMO1 -> CD32
SOD2 -> CD32
CD32 -> CD32
EIF -> CD32
G2/M -> CD32
S -> CD32
NA
RETN - TLR4
PTPRC - MRC1
NAMPT - INSR
MIF - (CD74+CXCR4)
MIF - (CD74+CXCR2)
MIF - (CD74+CD44)
LGALS9 - HAVCR2
LGALS9 - CD45
LGALS9 - CD44
ITGB2 - ICAM2
ITGB2 - ICAM1
ICAM2 - (ITGAM+ITGB2)
ICAM2 - (ITGAL+ITGB2)
ICAM1 - SPN
ICAM1 - ITGAL
ICAM1 - (ITGAX+ITGB2)
ICAM1 - (ITGAM+ITGB2)
ICAM1 - (ITGAL+ITGB2)
HLA-F - LILRB1
HLA-DRB5 - CD4
HLA-DRB1 - CD4
HLA-DRA - CD4
HLA-DQB1 - CD4
HLA-DQA2 - CD4
HLA-DQA1 - CD4
HLA-DPB1 - CD4
HLA-DPA1 - CD4
HLA-DMB - CD4
HLA-DMA - CD4
GRN - SORT1
GAS6 - AXL
FN1 - SDC4
FN1 - CD44
FN1 - (ITGAV+ITGB1)
FN1 - (ITGA5+ITGB1)
CD99 - PILRA
CD99 - CD99
APP - CD74
ANXA1 - FPR2
ANXA1 - FPR1
ADGRE5 - CD55
CD32 -> FN1
CD32 -> PLAC8
CD32 -> IFI27
CD32 -> CTNNB1
CD32 -> SCD
CD32 -> CTSC
CD32 -> MSMO1
CD32 -> SOD2
CD32 -> CD32
CD32 -> EIF
CD32 -> G2/M
CD32 -> S
Commun. Prob.
max
min
p-value
0.01 < p < 0.05
p < 0.01

J

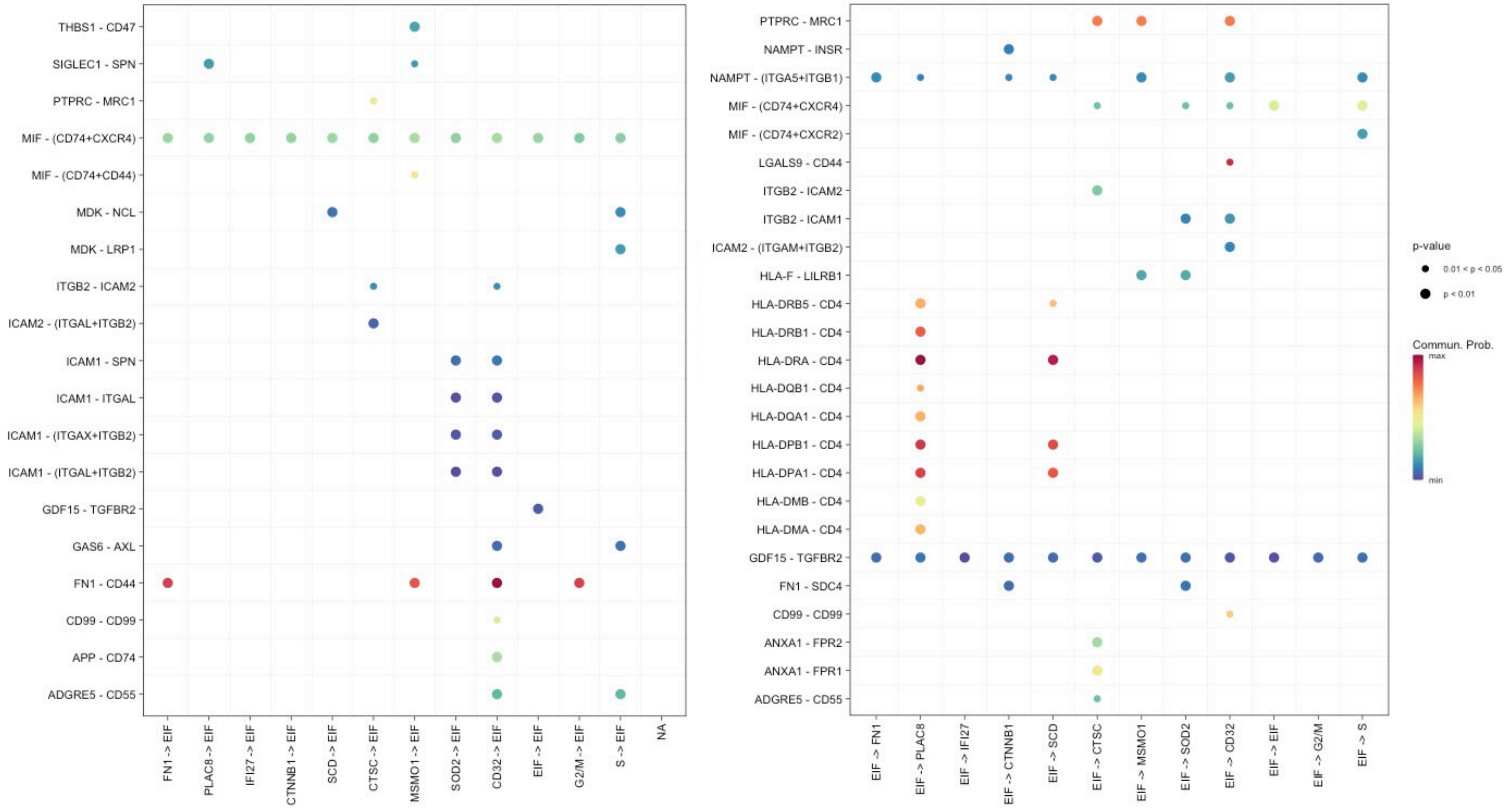
THBS1 - CD47
SIGLEC1 - SPN
PTPRC - MRC1
MIF - (CD74+CXCR4)
MIF - (CD74+CD44)
MDK - NCL
MDK - LRP1
ITGB2 - ICAM2
ICAM2 - (ITGAL+ITGB2)
ICAM1 - SPN
ICAM1 - ITGAL
ICAM1 - (ITGAX+ITGB2)
ICAM1 - (ITGAL+ITGB2)
GDF15 - TGFBR2
GAS6 - AXL
FN1 - CD44
CD99 - CD99
APP - CD74
ADGRE5 - CD55
FN1 -> EIF
PLAC8 -> EIF
IFI27 -> EIF
CTNNB1 -> EIF
SCD -> EIF
CTSC -> EIF
MSMO1 -> EIF
SOD2 -> EIF
CD32 -> EIF
EIF -> EIF
G2/M -> EIF
S -> EIF
NA
PTPRC - MRC1
NAMPT - INSR
NAMPT - (ITGA5+ITGB1)
MIF - (CD74+CXCR4)
MIF - (CD74+CXCR2)
LGALS9 - CD44
ITGB2 - ICAM2
ITGB2 - ICAM1
ICAM2 - (ITGAM+ITGB2)
HLA-F - LILRB1
HLA-DRB5 - CD4
HLA-DRB1 - CD4
HLA-DRA - CD4
HLA-DQB1 - CD4
HLA-DQA1 - CD4
HLA-DPB1 - CD4
HLA-DPA1 - CD4
HLA-DMB - CD4
HLA-DMA - CD4
GDF15 - TGFBR2
FN1 - SDC4
CD99 - CD99
ANXA1 - FPR2
ANXA1 - FPR1
ADGRE5 - CD55
EIF -> FN1
EIF -> PLAC8
EIF -> IFI27
EIF -> CTNNB1
EIF -> SCD
EIF -> CTSC
EIF -> MSMO1
EIF -> SOD2
EIF -> CD32
EIF -> EIF
EIF -> G2/M
EIF -> S
p-value
0.01 < p < 0.05
p < 0.01
Commun. Prob.
max
min

K

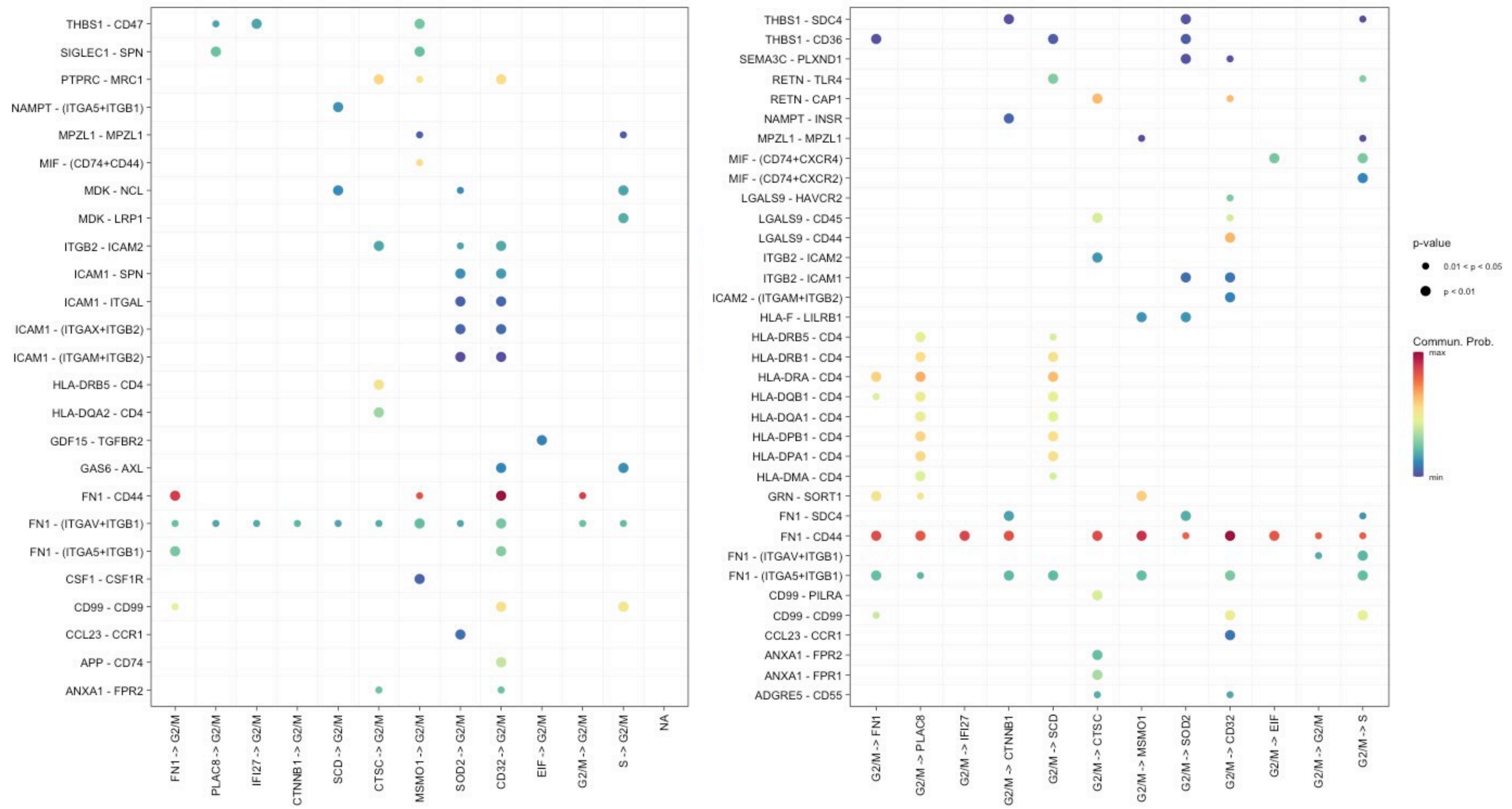
THBS1 - CD47
SIGLEC1 - SPN
PTPRC - MRC1
NAMPT - (ITGA5+ITGB1)
MPZL1 - MPZL1
MIF - (CD74+CD44)
MDK - NCL
MDK - LRP1
ITGB2 - ICAM2
ICAM1 - SPN
ICAM1 - ITGAL
ICAM1 - (ITGAX+ITGB2)
ICAM1 - (ITGAM+ITGB2)
HLA-DRB5 - CD4
HLA-DQA2 - CD4
GDF15 - TGFBR2
GAS6 - AXL
FN1 - CD44
FN1 - (ITGAV+ITGB1)
FN1 - (ITGA5+ITGB1)
CSF1 - CSF1R
CD99 - CD99
CCL23 - CCR1
APP - CD74
ANXA1 - FPR2
FN1 -> G2/M
PLAC8 -> G2/M
IFI27 -> G2/M
CTNNB1 -> G2/M
SCD -> G2/M
CTSC -> G2/M
MSMO1 -> G2/M
SOD2 -> G2/M
CD32 -> G2/M
EIF -> G2/M
G2/M -> G2/M
S -> G2/M
NA
THBS1 - SDC4
THBS1 - CD36
SEMA3C - PLXND1
RETN - TLR4
RETN - CAP1
NAMPT - INSR
MPZL1 - MPZL1
MIF - (CD74+CXCR4)
MIF - (CD74+CXCR2)
LGALS9 - HAVCR2
LGALS9 - CD45
LGALS9 - CD44
ITGB2 - ICAM2
ITGB2 - ICAM1
ICAM2 - (ITGAM+ITGB2)
HLA-F - LILRB1
HLA-DRB5 - CD4
HLA-DRB1 - CD4
HLA-DRA - CD4
HLA-DQB1 - CD4
HLA-DQA1 - CD4
HLA-DPB1 - CD4
HLA-DPA1 - CD4
HLA-DMA - CD4
GRN - SORT1
FN1 - SDC4
FN1 - CD44
FN1 - (ITGAV+ITGB1)
FN1 - (ITGA5+ITGB1)
CD99 - PILRA
CD99 - CD99
CCL23 - CCR1
ANXA1 - FPR2
ANXA1 - FPR1
ADGRE5 - CD55
G2/M -> FN1
G2/M -> PLAC8
G2/M -> IFI27
G2/M -> CTNNB1
G2/M -> SCD
G2/M -> CTSC
G2/M -> MSMO1
G2/M -> SOD2
G2/M -> CD32
G2/M -> EIF
G2/M -> G2/M
G2/M -> S
p-value
0.01 < p < 0.05
p < 0.01
Commun. Prob.
max
min

L

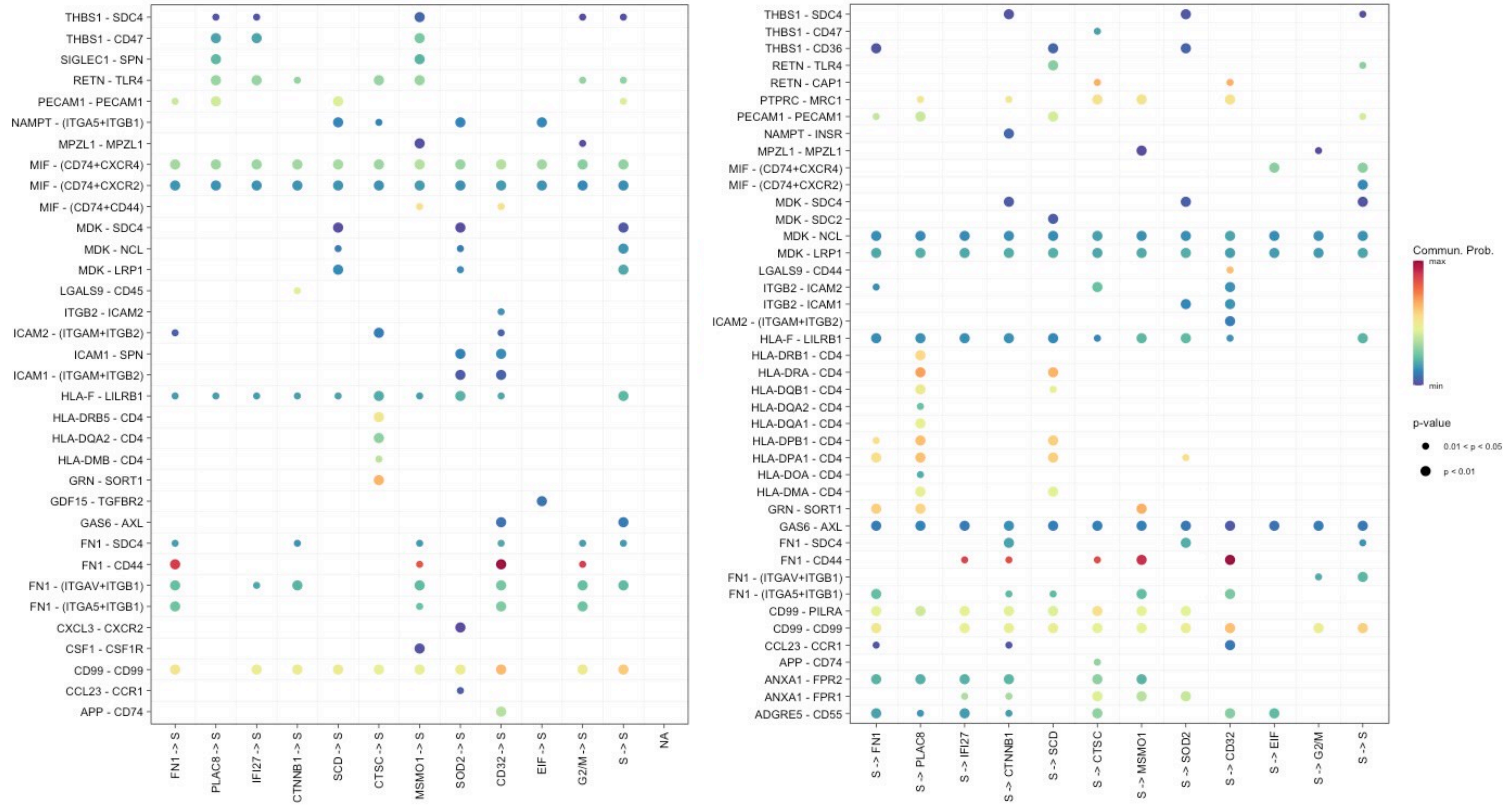
THBS1 - SDC4
THBS1 - CD47
SIGLEC1 - SPN
RETN - TLR4
PECAM1 - PECAM1
NAMPT - (ITGA5+ITGB1)
MPZL1 - MPZL1
MIF - (CD74+CXCR4)
MIF - (CD74+CXCR2)
MIF - (CD74+CD44)
MDK - SDC4
MDK - NCL
MDK - LRP1
LGALS9 - CD45
ITGB2 - ICAM2
ICAM2 - (ITGAM+ITGB2)
ICAM1 - SPN
ICAM1 - (ITGAM+ITGB2)
HLA-F - LILRB1
HLA-DRB5 - CD4
HLA-DQA2 - CD4
HLA-DMB - CD4
GRN - SORT1
GDF15 - TGFBR2
GAS6 - AXL
FN1 - SDC4
FN1 - CD44
FN1 - (ITGAV+ITGB1)
FN1 - (ITGA5+ITGB1)
CXCL3 - CXCR2
CSF1 - CSF1R
CD99 - CD99
CCL23 - CCR1
APP - CD74
FN1 -> S
PLAC8 -> S
IFI27 -> S
CTNNB1 -> S
SCD -> S
CTSC -> S
MSMO1 -> S
SOD2 -> S
CD32 -> S
EIF -> S
G2/M -> S
S -> S
NA
THBS1 - SDC4
THBS1 - CD47
THBS1 - CD36
RETN - TLR4
RETN - CAP1
PTPRC - MRC1
PECAM1 - PECAM1
NAMPT - INSR
MPZL1 - MPZL1
MIF - (CD74+CXCR4)
MIF - (CD74+CXCR2)
MDK - SDC4
MDK - SDC2
MDK - NCL
MDK - LRP1
LGALS9 - CD44
ITGB2 - ICAM2
ITGB2 - ICAM1
ICAM2 - (ITGAM+ITGB2)
HLA-F - LILRB1
HLA-DRB1 - CD4
HLA-DRA - CD4
HLA-DQB1 - CD4
HLA-DQA2 - CD4
HLA-DQA1 - CD4
HLA-DPB1 - CD4
HLA-DPA1 - CD4
HLA-DOA - CD4
HLA-DMA - CD4
GRN - SORT1
GAS6 - AXL
FN1 - SDC4
FN1 - CD44
FN1 - (ITGAV+ITGB1)
FN1 - (ITGA5+ITGB1)
CD99 - PILRA
CD99 - CD99
CCL23 - CCR1
APP - CD74
ANXA1 - FPR2
ANXA1 - FPR1
ADGRE5 - CD55
S -> FN1
S -> PLAC8
S -> IFI27
S -> CTNNB1
S -> SCD
S -> CTSC
S -> MSMO1
S -> SOD2
S -> CD32
S -> EIF
S -> G2/M
S -> S
Commun. Prob.
max
min
p-value
0.01 < p < 0.05
p < 0.01

Figure S7

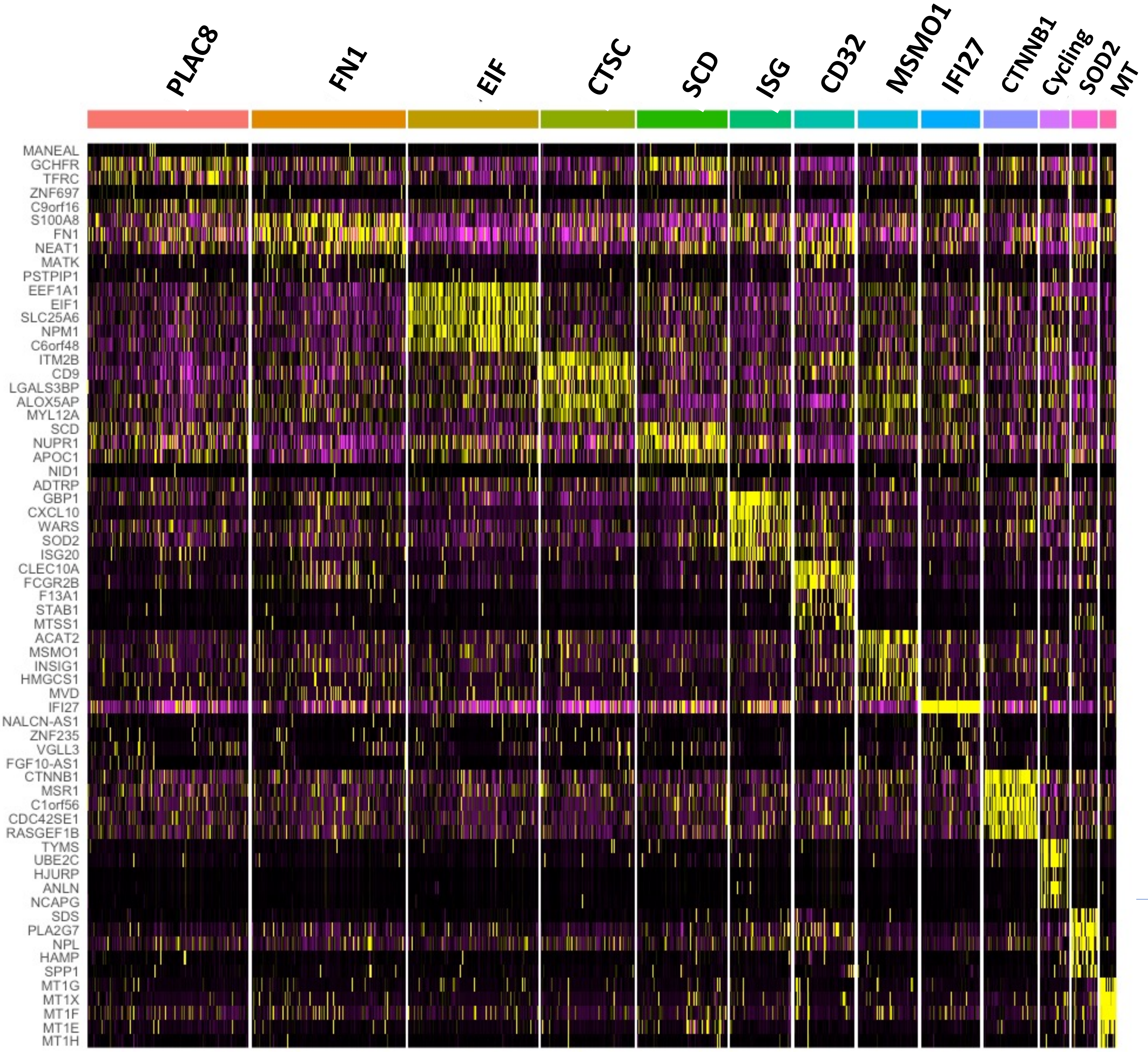
PLAC8
FN1
EIF
CTSC
SCD
ISG
CD32
MSMO1
IFI27
CTNNB1
Cycling
SOD2
MT
MANEAL
GCHFR
TFRC
ZNF697
C9orf16
S100A8
FN1
NEAT1
MATK
PSTPIP1
EEF1A1
EIF1
SLC25A6
NPM1
C6orf48
ITM2B
CD9
LGALS3BP
ALOX5AP
MYL12A
SCD
NUPR1
APOC1
NID1
ADTRP
GBP1
CXCL10
WARS
SOD2
ISG20
CLEC10A
FCGR2B
F13A1
STAB1
MTSS1
ACAT2
MSMO1
INSIG1
HMGCS1
MVD
IFI27
NALCN-AS1
ZNF235
VGLL3
FGF10-AS1
CTNNB1
MSR1
C1orf56
CDC42SE1
RASGEF1B
TYMS
UBE2C
HJURP
ANLN
NCAPG
SDS
PLA2G7
NPL
HAMP
SPP1
MT1G
MT1X
MT1F
MT1E
MT1H

**Figure S8**

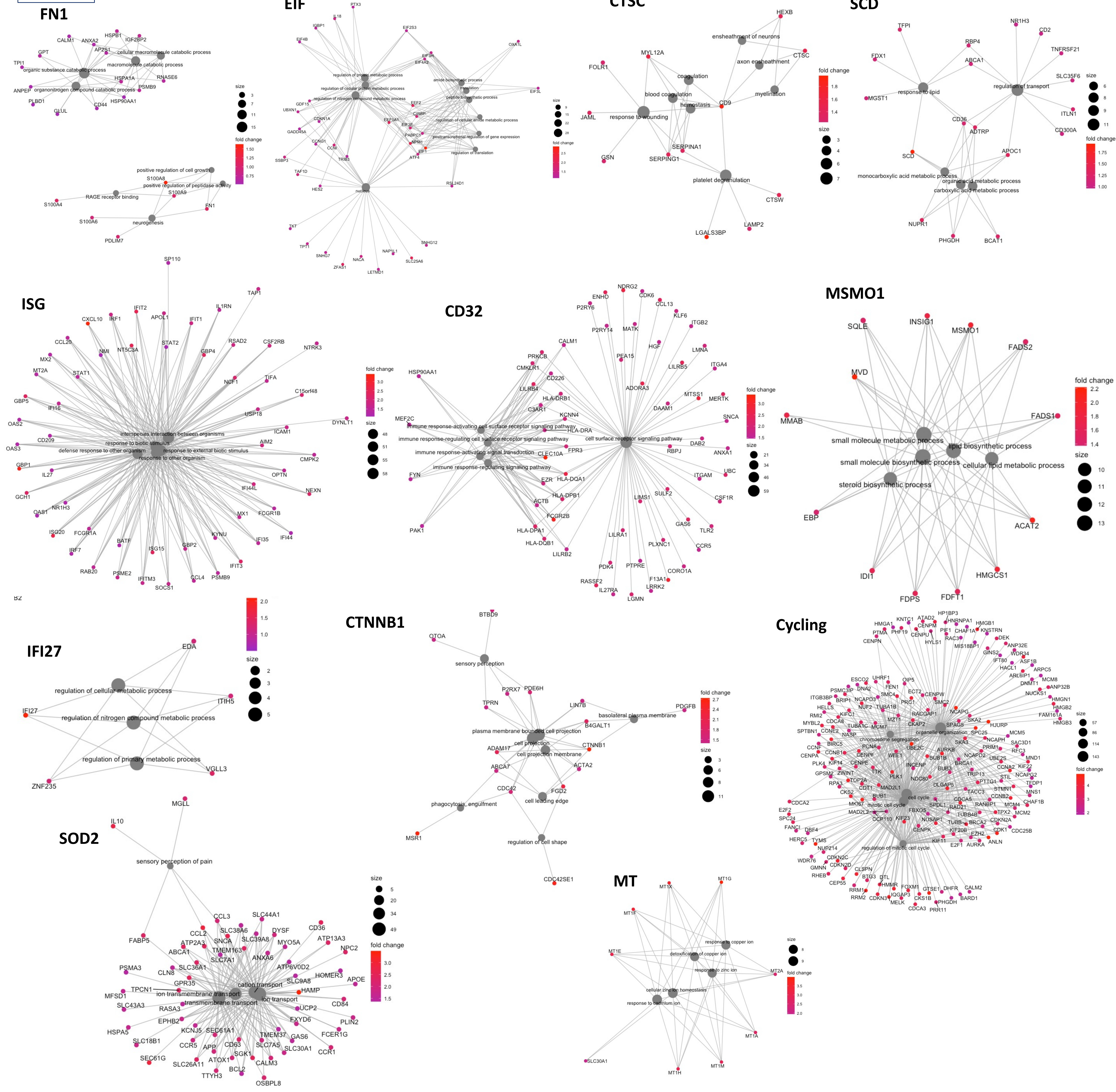

FN1
cellular macromolecule catabolic process
macromolecule catabolic process
organic substance catabolic process
organonitrogen compound catabolic process
positive regulation of cell growth
positive regulation of peptidase activity
RAGE receptor binding
neurogenesis
fold change
size
EIF
CTSC
ensheathment of neurons
axon ensheathment
coagulation
blood coagulation
hemostasis
myelination
response to wounding
platelet degranulation
SCD
response to lipid
regulation of transport
monocarboxylic acid metabolic process
organic acid metabolic process
carboxylic acid metabolic process
ISG
interspecies interaction between organisms
response to biotic stimulus
defense response to other organism
response to external biotic stimulus
response to other organism
CD32
immune response-activating cell surface receptor signaling pathway
immune response-regulating cell surface receptor signaling pathway
immune response-activating signal transduction
immune response-regulating signaling pathway
cell surface receptor signaling pathway
MSMO1
small molecule metabolic process
lipid biosynthetic process
small molecule biosynthetic process
cellular lipid metabolic process
steroid biosynthetic process
IFI27
regulation of cellular metabolic process
regulation of nitrogen compound metabolic process
regulation of primary metabolic process
CTNNB1
sensory perception
basolateral plasma membrane
plasma membrane bounded cell projection
cell projection
cell projection membrane
phagocytosis, engulfment
cell leading edge
regulation of cell shape
Cycling
SOD2
sensory perception of pain
cation transport
ion transmembrane transport
transmembrane transport
ion transport
MT
response to copper ion
detoxification of copper ion
response to zinc ion
cellular zinc ion homeostasis
response to cadmium ion

Figure S9

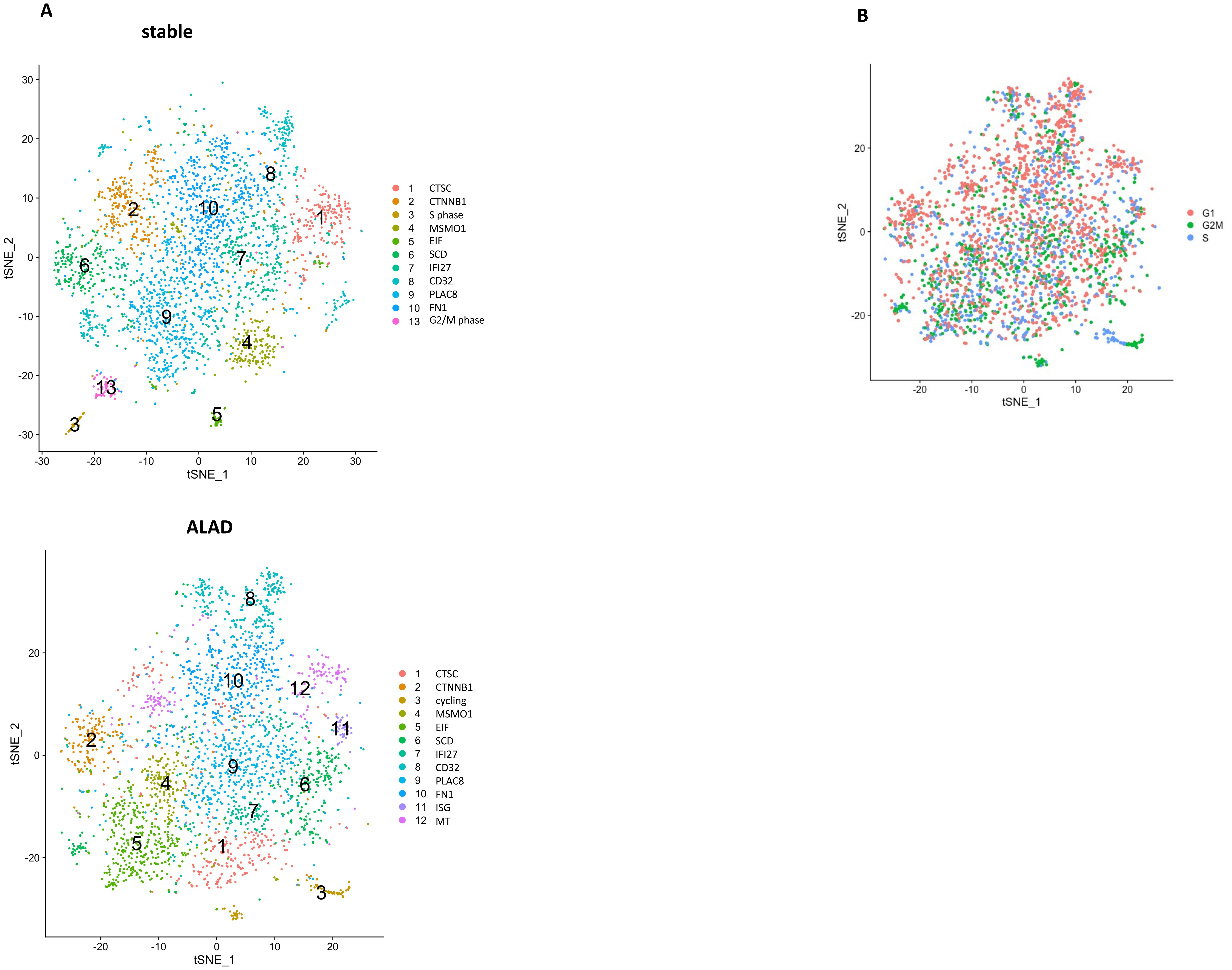

A
stable
30
20
10
0
-10
-20
-30
tSNE_2
tSNE_1
1
2
3
4
5
6
7
8
9
10
13
1 CTSC
2 CTNNB1
3 S phase
4 MSMO1
5 EIF
6 SCD
7 IFI27
8 CD32
9 PLAC8
10 FN1
13 G2/M phase
B
tSNE_2
tSNE_1
G1
G2M
S
ALAD
tSNE_2
tSNE_1
1 CTSC
2 CTNNB1
3 cycling
4 MSMO1
5 EIF
6 SCD
7 IFI27
8 CD32
9 PLAC8
10 FN1
11 ISG
12 MT

**Figure S10**

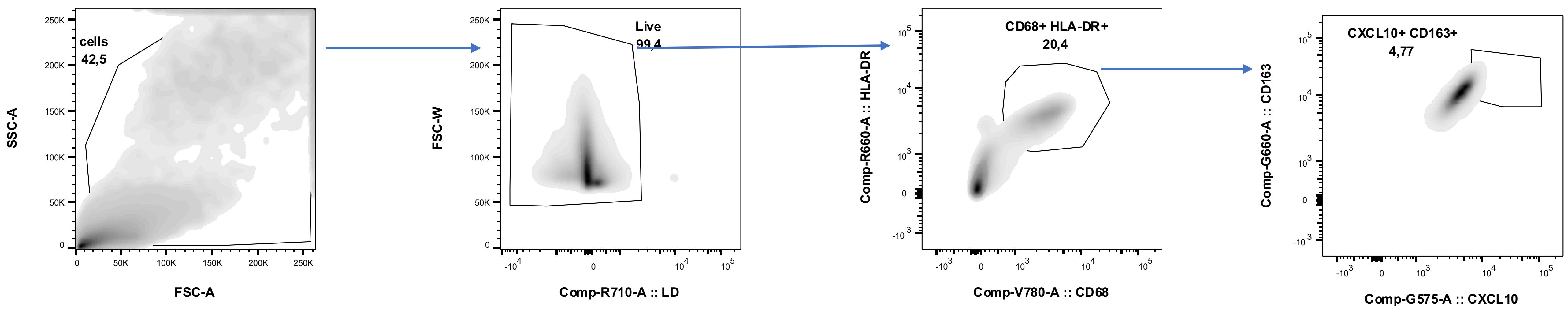
cells
42,5
SSC-A
FSC-A
Live
99,4
FSC-W
Comp-R710-A :: LD
CD68+ HLA-DR+
20,4
Comp-R660-A :: HLA-DR
Comp-V780-A :: CD68
CXCL10+ CD163+
4,77
Comp-G660-A :: CD163
Comp-G575-A :: CXCL10

Figure S11

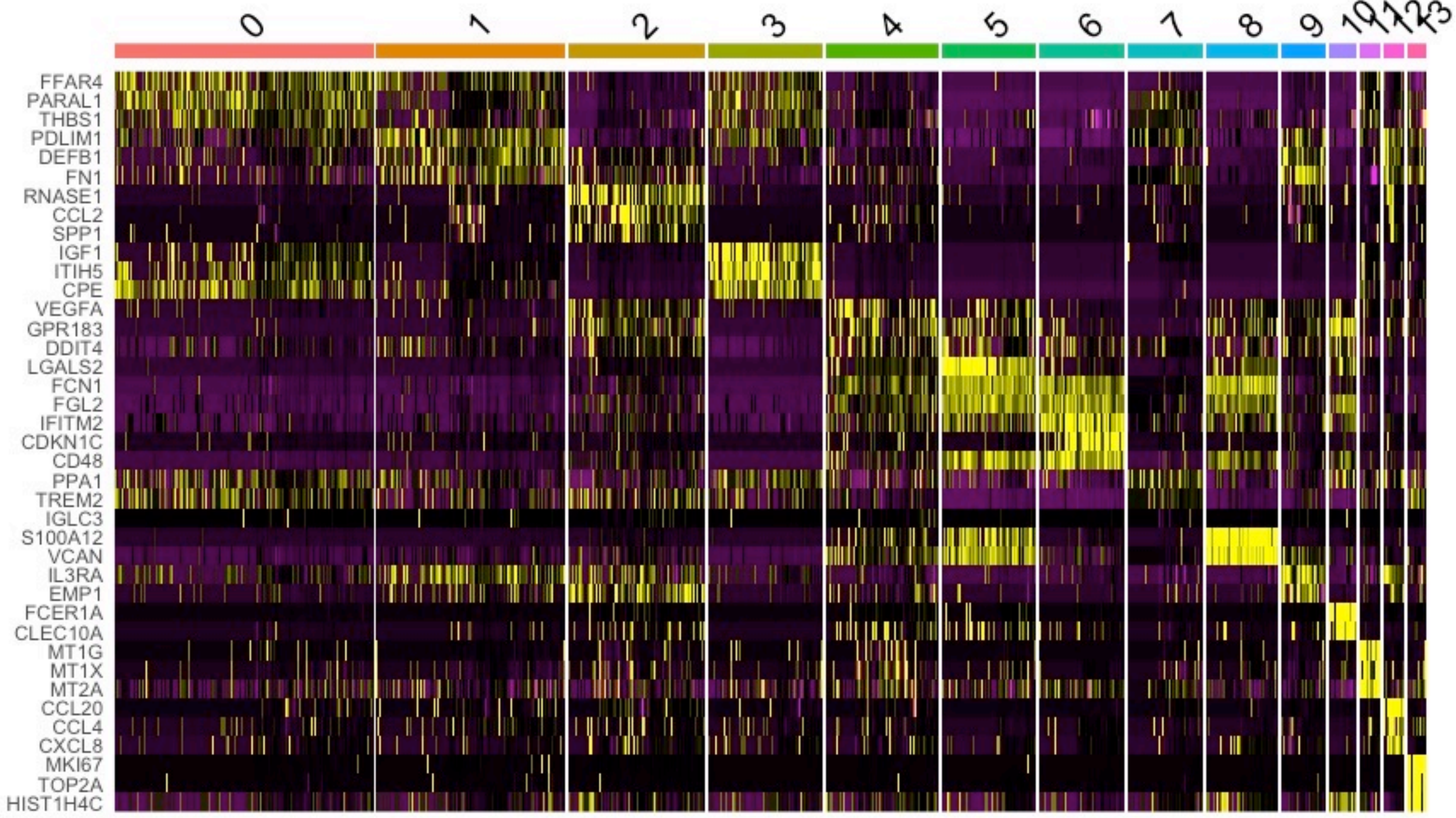
0
1
2
3
4
5
6
7
8
9
10
11
12
13
FFAR4
PARAL1
THBS1
PDLIM1
DEFB1
FN1
RNASE1
CCL2
SPP1
IGF1
ITIH5
CPE
VEGFA
GPR183
DDIT4
LGALS2
FCN1
FGL2
IFITM2
CDKN1C
CD48
PPA1
TREM2
IGLC3
S100A12
VCAN
IL3RA
EMP1
FCER1A
CLEC10A
MT1G
MT1X
MT2A
CCL20
CCL4
CXCL8
MKI67
TOP2A
HIST1H4C
